# Personalized cancer treatment strategies incorporating irreversible and reversible drug resistance mechanisms

**DOI:** 10.1101/2024.11.03.621749

**Authors:** Wei He, Matthew D. McCoy, Rebecca B. Riggins, Robert A. Beckman, Chen-Hsiang Yeang

## Abstract

Despite advances in targeted cancer therapy, the promise of precision medicine has been limited by resistance to these treatments. In this study, we propose a mathematical modelling framework incorporating cellular heterogeneity, genetic evolutionary dynamics, and non-genetic plasticity, accounting for both irreversible and reversible drug resistance. Previously we proposed Dynamic Precision Medicine (DPM), a personalized treatment strategy that designed individualized treatment sequences by simulations of irreversible genetic evolutionary dynamics in a heterogeneous tumor. Here we apply DPM to the joint model of reversible and irreversible drug resistance mechanisms, analyze the simulation results and compare the efficacy of various treatment strategies. The results indicate that this enhanced version of DPM results in superior patient outcomes compared with current personalized medicine treatment approaches. Our results provide insights into cancer treatment strategies for heterogeneous tumors with genetic evolutionary dynamics and non-genetic cellular plasticity, potentially leading to improvements in survival time for cancer patients.

## INTRODUCTION

Drug resistance remains a primary obstacle in cancer treatment^1^. Intratumoral genetic heterogeneity and non-genetic plasticity in cancer cells are two major factors of cancer treatment resistance^2–9^, and are widely associated with poor outcomes and reduced responses to therapies^1,2,10–13^. A bulk tumor typically consists of a heterogeneous population of cancer cells often characterized by bulk or subclonal genetic instability^2,10,14–17^. By sequencing tumors with a higher depth and accuracy and applying a novel mathematical model, we found that every DNA nucleotide is mutated in at least one cell in the smallest tumor visible on CT (Computed Tomography), unless that mutation is highly selected against^14^. This implies that at least one cancer cell is likely resistant to any single drug therapy and already present in any tumor at diagnosis^14^. Elevated genetic mutation rates and selection imposed by treatments drive the clonal evolution of tumors toward resistance. Of note, it is likely that a subset of cells has mutations in the cellular machinery that normally ensures genetic stability. We have termed these subclones “hypermutator subclones,” and they may be selected as single and multidrug resistance evolves^17^. In addition, non-genetic plasticity in cancer cells provides a rapid reversible mechanism for drug-induced resistance^13^. Cancer cells can increase drug efflux, alter drug metabolism and activate bypassing signaling pathways to become resistant to the drugs. These two types of drug resistance mechanisms – intratumoral genetic subclonal heterogeneity and non-genetic cellular plasticity and heterogeneity – differ in their reversibility. The former is irreversible as a resistant subclone will rarely revert mutations to lose its phenotype before expanding, and then reversions will occur in only a minority of cells^18^. It is likely to cause moderate term to late progression or relapse as it may involve outgrowth of rare subclones, and/or the accumulation of multiple resistance mutations. The latter is reversible as cells can alter their internal states to adapt to changes in the microenvironment (e.g., presence or absence of drugs)^13,19,20^, hence resistance can be reversed when the drug treatment is discontinued^20^. This reversible resistance may occur rapidly in a majority of cells, and clinically may correspond to primary resistance and/or short term relapses.

Previously, we have proposed mathematical models and treatment strategies to address the irreversible and reversible mechanisms of drug resistance in separate studies^21–23^. For the irreversible mechanism, we constructed a model to capture the population dynamics of tumor subclones as they acquire resistance to two non-cross resistant drugs through independent mutations, and proposed a treatment selection strategy, Dynamic Precision Medicine (DPM) to design the treatment sequence to balance the immediate goal of shrinking tumor size and the long term goal of preventing the emergence of an incurable subclone resistant to both drugs^21^. (Notably, many drugs are cross resistant. Such drugs are considered the same drug by DPM, and even when used in combination are treated as a single “drug”). DPM contrasts with the conventional precision medicine approach that attempts to match a drug (or a combination of drugs) to the molecular profile of a patient but does not address the complex relations between the patient’s molecular profile, possible treatment sequences, and the dynamic response of the tumor. We further extended the model to include three drugs and incorporated long-term prediction of tumor population dynamics^22^. For the reversible mechanism, we introduced another mathematical model where cancer cells responded to the presence or absence of a drug by activating/inhibiting two alternative pathways, and proposed an optimal dynamic treatment strategy accordingly^23^.

We now report a single, integrated mathematical model that captures both irreversible genetic drug resistance and reversible drug resistance induced by cellular plasticity. The unified framework encompasses both irreversible and reversible drug resistance for two non-cross resistant drugs, and the treatment strategies that simultaneously tackle the irreversible and reversible drug resistance mechanisms. We evaluate the effectiveness of nine treatment strategies by simulating the dynamics of cancer cell populations. We conduct a clinical trial simulation over 6 million virtual patients, each representing a different dynamic presentation at diagnosis, and demonstrate that the DPM-based personalized treatment strategies result in superior patient outcomes compared with the current personalized medicine treatment approach. Furthermore, DPM strategies incorporating periodic treatment sequences that cycle between therapies over a shorter treatment window, designed to combat reversible resistance, are marginally superior to those without such options.

Mathematical models have been utilized to design effective treatment strategies aimed at addressing drug resistance^24–29^. Gatenby and colleagues proposed adaptive therapy, a strategy that dynamically adjusts treatment to preserve a population of drug-sensitive cells. These cells are proposed to compete with resistant ones, with the goal of controlling the tumor rather than fully eradicating it^30^. However, these sensitive cells continue to mutate and serve as a reservoir for future variants, and the competition between cells may be less of a factor in small metastatic sites that are usually the primary cause of death in patients^31^. Moreover, sensitive cells have in some cases been documented to support the fitness of resistant cells in co-culture^32,33^. Strobl et al. proposed that adaptive therapy could mitigate both the toxicity and resistance associated with poly-adenosine ribose polymerase inhibitor (PARPi)^26^. Gallagher et al. applied deep reinforcement learning to guide adaptive drug scheduling, demonstrating that such schedules could outperform the current adaptive protocols in a mathematical model calibrated to prostate cancer, helping to delay the onset of drug resistance. They tested their mathematical models on a limited number of virtual patients, whereas we examined our model through simulations involving a virtual clinical trial of over 6 million patients. Adaptive therapy has shown promising results in metastatic prostate cancer treating with a single drug^34^. However, sensitive and resistant cells were inferred in this model rather than measured, and it is possible that the therapy interruptions in adaptive therapy with a single agent are effective because they essentially promote a cycling approach, which can also be beneficial for managing reversible resistance, rather than due to competitive interactions between sensitive and resistant cells. Our approach aims to delay or prevent the appearance of doubly genetically resistant cell types and was one of the first to include sequencing of multiple non-cross resistant therapies by using multiple agents^21^. Cárdenas et al. proposed a family of mathematical models to investigate the timing and mechanism leading to irreversible resistance to cetuximab in head and neck squamous cell carcinoma^11^. Sahoo et al. explored the coupled dynamics of epithelial-mesenchymal transition (EMT) and the development of reversible drug resistance in breast cancer^35^. Their findings suggest a reciprocal relationship between EMT and resistance to tamoxifen, where each process can promote the other. This offers a mechanistic explanation for empirical observations that cells undergoing EMT tend to be more resistant to tamoxifen and that tamoxifen-resistant cells often exhibit EMT-like characteristics. These models typically focus on a single resistance mechanism, either reversible or irreversible. In contrast, we propose a mathematical modelling framework incorporating both irreversible and reversible drug resistance. This approach enhances existing models by providing insights into treatment strategies for heterogeneous tumors, accounting for both Intratumoral genetic heterogeneity and non-genetic cellular plasticity.

## METHODS

### Model Overview

We developed a joint mathematical model that incorporates both reversible and irreversible resistance mechanisms to simulate virtual patients’ outcomes under different treatment strategies. We give a high-level overview of the model here and provide more precise descriptions in the subsequent sections. The irreversible and reversible portions of the model are primarily based on our prior studies^21–23^. The model includes two drugs, drug 1 and drug 2. In the irreversible part of the model, response of each drug in a cancer cell is indicated by a binary phenotype (0 denotes sensitive and 1 resistant responses), and drug resistance is acquired by the mutation of a gene (or a group of genes). There are four phenotypes in combination and we represent them by two bits: 00, 01, 10, 11, representing the phenotypes sensitive to both drugs, resistant to drug 2, drug 1, and both drugs, respectively. These four cell types manifest differential death rates induced by drug dosages, and only mutations from sensitive to resistant phenotypes are allowed. In the reversible part of the model, net growth rate (proliferation rate minus death rate) of a cancer cell can be increased by two distinct pathways, and the activity of each pathway in a cancer cell is again indicated by a binary phenotype (0 representing an inactive and 1 an active state). There are also four cellular states in combination and we represent them by two bits: 00, 01, 10, 11 denote the activities of two pathways. The two drugs target the two pathways by inhibiting their activities. Therefore, the dosage of a drug affects population dynamics in two ways: (1) reducing the net growth rate of the cells with high activity of the targeted pathway, (2) facilitating cellular state transitions from active to inactive targeted pathway. Also, cells with an inactive pathway are resistant to the drug targeting the pathway. The irreversible drug resistance phenotypes and reversible pathway activities together define four-bit state vectors. Bits 1 and 3 denote the irreversible resistance phenotypes of drugs 1 and 2, and bits 2 and 4 denote the reversible activity states of the targeted pathways of drugs 1 and 2.

In total there are nine stable cell types representing distinct phenotypic states, because certain states contradict with the definitions hence will quickly transition into stable states. For instance, 0010 is transient since it has an irreversible resistance to drug 2 but inactive pathway 2 implicating potential drug 2 action. The irreversible resistance drives activation of the proliferation pathway and it will quickly transition to the 0011 state.

The description of these nine stable states is shown in Table 1. In the model with only reversible resistance, bits 1 and 3 are 0s and there are only four valid states: 0101, 0001, 0100 and 0000. Similarly, in the model with only irreversible resistance, bits 2 and 4 are 1s and there are only four stable states: 0101, 1101, 0111 and 1111, where 0101 is a valid state in both irreversible and reversible models. There are two additional states, 1100 and 0011, that emerge when integrating the two resistance mechanisms. There are transient states-1000, 1001, 1110, 1011, 0010, and 0110, and -which will quickly transition to 1100, 1101, 1111, 1111, 0011, and 0111, respectively, as shown in Fig. 1a. The 1010 state is unlikely to occur in that it postulates two independent irreversible resistance mutations arising simultaneously in the same cell, if the drugs are non-cross resistant. Rather, it is more likely that one resistance mutation would arise first, and its corresponding reversible state would rapidly revert to 1 if it were 0. In total there are 16 states, 6 of which are transient. The transient states are therefore omitted in the joint model, as illustrated in Fig. 1b. 0101 represents the predominant cell type identified in the biopsy results when a cancer patient is first diagnosed. In our model, drug 1 corresponds to the first line treatment for this predominant cell type. Therefore, drug 1 is considered more effective against 0101 cells than drug 2.

**Table 1.** Cell state descriptions.

| Index | Cell state | Description |
| --- | --- | --- |
| 1 | 0101 | sensitive to drug 1 and drug 2 |
| 2 | 0001 | reversible resistant to drug 1 and sensitive to drug 2 |
| 3 | 0100 | sensitive to drug 1 and reversible resistant to drug 2 |
| 4 | 0000 | reversible resistant to both drug 1 and drug 2 |
| 5 | 0111 | sensitive to drug 1 and irreversible resistant to drug 2 |
| 6 | 0011 | reversible resistant to drug 1 and irreversible resistant to drug 2 |
| 7 | 1101 | irreversible resistant to drug 1 and sensitive to drug 2 |
| 8 | 1100 | irreversible resistant to drug 1 and reversible resistant to drug 2 |
| 9 | 1111 | irreversible resistant to both drug 1 and drug 2 |

**Fig 1:**
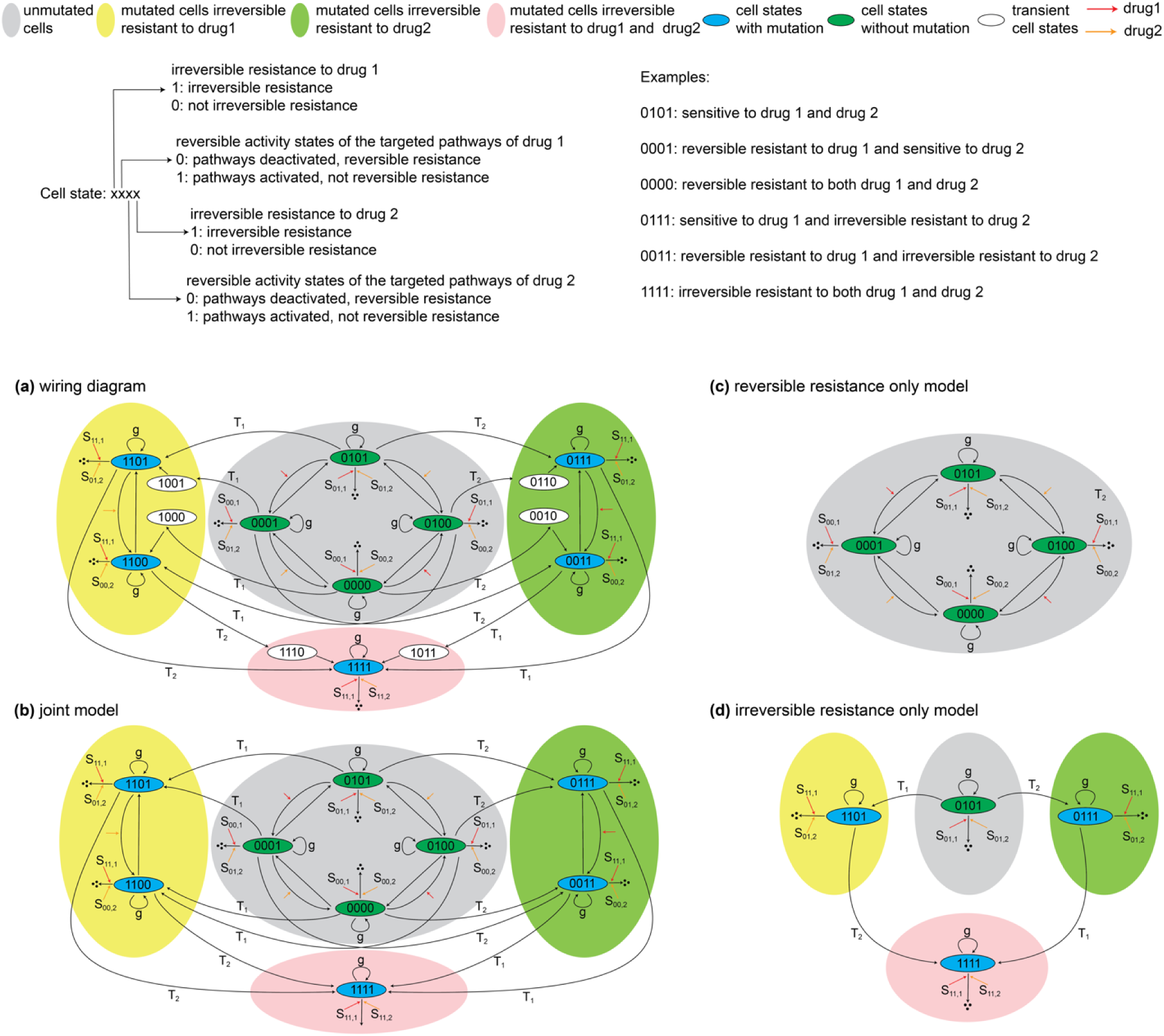
Overview of proliferations and transitions among different cell states and schematic representation of the structures of the mathematical model. **a:** The schematic illustrating the dynamics of different cell states and their transitions in the presence of drug 1 and drug 2. Cell states are represented by 4 bits. Bits 1 and 3 denote the irreversible resistance phenotypes of drugs 1 and 2, and bits 2 and 4 denote the reversible activity states of the targeted pathways of drugs 1 and 2. The central gray bubble comprises the state transitions of the reversible resistance only model, i.e., bits 1 and 3 are 0 or the irreversible resistance mutations do not occur. The left yellow bubble and the right green bubble comprise the state transitions where the irreversible resistance mutation of drug 1 (bit 1) or drug 2 (bit 3) alone occurs. The white states (1001, 1000, 0110, 0010, 1110, 1011) represent transient states where mutations have occurred but pathways have not been activated. These transient states will quickly transition to 1101, 1100, 0111 and 0011, respectively. The bottom pink bubble comprises the states possessing double irreversible resistance of both drugs, including one stable (1111) and two transient (1110, 1011) states. Arrows include self-replications, degradations and transitions/mutations from or to designated states. **b:** The joint model structure used for simulation. Transient states are omitted compared to **a**. **c:** the reversible resistance only model. **d:** the irreversible resistance only model.

The interplay between the internal states of cancer cells and the external conditions exerted by the drug dosages has the potential to produce a variety of population dynamics depending on the sequencing or dosage. Population dynamics are driven by both long-term irreversible and short-term reversible drug resistance mechanisms. In the long term, sensitive and resistant phenotypes manifest differential net growth rates in response to the administered drugs, and transition rates from sensitive to irreversible resistant phenotypes are independent of the drug dosages (dependencies on drug dosages can be introduced when considering drugs that interfere with DNA replication and repair, but were not considered in the current work). The genetic transitions of these abstract phenotypes can be attributed to all possible genetic mutations that induce the irreversible resistance phenotype. Drug dosages affect population dynamics by eliminating (or reducing) the sensitive subclones hence selecting the resistant subclones. In the short term, net growth rate is enhanced by either one of two pathways. Drug treatment inhibits the targeted pathway activity and kills the sensitive cells, but it also facilitates the activation of alternative pathways, leading to transitions that induce drug resistance. When a drug treatment is withdrawn, the targeted pathway is no longer inhibited, hence the cells likely transition back to the sensitive state. The transition rates between these reversible states thereby depend on the drug dosage. A high dosage facilitates transitions from sensitive to resistant states, while a low (or zero) dosage facilitates transitions of the opposite direction.

### Model Parameters

Supplementary Table 1 lists the 17 parameters in the joint model and their value ranges in simulations. They belong to five general categories. (1) g represents the natural net growth rate which, in these simulations is assumed to be identical for all cell types. This will not generally be the case, and separate natural net growth rates can be input into the simulations if known, resulting in results that are applicable to those cases. The term “natural” refers to the rate in absence of drug therapy. (2) α_1_, θ_1_, μ_1_, α_2_, θ_2_, μ_2_ represent the parameters pertaining to the transitions between the reversible states governed by the activities of the two pathways. Their meanings will be explained in the next section. (3) T_1_ and T_2_ represent the rates of acquiring irreversible resistance phenotypes to each drug by genetic change. (4) S_01,1_ is drug 1 sensitivity of sensitive cells relative to the natural net growth rate; S_01,2_ is drug 2 sensitivity of sensitive cells relative to the net natural growth rate ; S_00,1_, is drug 1 sensitivity relative to the natural net growth rate for cells reversibly resistant to drug 1; S_00,2_, is drug 2 sensitivity relative to the net natural growth rate for cells reversibly resistant to drug 1; S_11,1_, is drug 1 sensitivity relative to net natural growth rate for cells irreversibly resistant to drug 1; S_11,2_ is drug 2 sensitivity relative to natural net growth rate for cells irreversibly resistant to drug 2. The first two indices indicate the irreversible resistance phenotype (bit 1) and the pathway activity (bit 2), and the third index indicates the targeted phenotype/pathway of each drug (1 or 2). (5) R_1ratio_ and R_2ratio_ represent the fractions of cells possessing irreversible resistance of drugs 1 and 2 in the initial population. We set the cells with double irreversible resistance to be absent (n1111=0) in the initial population for all simulations since all treatments are ineffective against those cells.

Cell types possessing reversible resistance alone (S_00,x_, x=1 or 2) are less sensitive to the targeted drug compared to cell types without resistance (S_01,x_) where S_00,x_= S_01,x_ × SR_00,x_ and SR_00,x_<1. Likewise, cell types possessing irreversible resistance alone (S_11,x_) are less sensitive to the targeted drug compared to cell types without resistance (S_01,x_), where S_11,x_= S_01,x_ × SR_11,x_ and SR_11,x_<1. We also set SR_11,x_ < SR_00,x_, indicating that irreversible resistance leads to a more robust and persistent effect compared to reversible resistance. Furthermore, as a convention we assume drug 1 is more effective than drug 2 in eliminating sensitive cells, hence set S_01,1 ≥_ S_01,2_. The model parameter value ranges are based on clinical, in vitro and in vivo data^21^. By including possible parameter values, we can state general conclusions across a broad range of individual virtual patients.

### Modelling equations

The transition rates between sensitive and reversibly resistant cell states are defined by parameters α_i_, θ_i_, μ_i_, i=1, 2 as in our previous work^23^. We first briefly recapitulate the reversible part of the model. Denote *y_{_* pathway *i* activity. The dynamics of *y_i_*(*t*) follow a first-order differential equation with production function *f(y_i_,σ_i_,θ_i_,α_i_)*, degradation term proportional to *y_i_(t)*, relative adjustment rate *δ* and random noise term *η*_i_

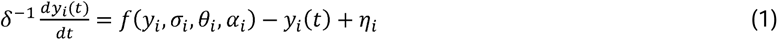

*f(y_i_,σ_i_,θ_i_,α_i_)* depends on drug *i* dosage *σ_i_* and *parameters α_i_, θ_i_.* We set 0 ≤ *α_i_*, ≤ *θ_i_* ≤ 1 to make the production function is a step-wise function^23^, it is proportional to one for any *y_i_* above threshold *θ_i_* and to *α_i_* for any *y_i_* below it. *f(y_i_,σ_i_,θ_i_,α_i_)* has the following form:

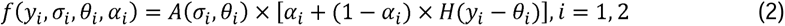

where *A*(*σ_i_,θ_i_*) is a decreasing function of *σ_i_*.

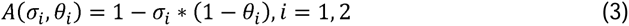

and *H(°)* is the Heaviside (step) function:

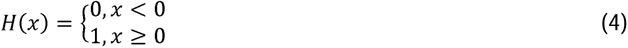

Qualitatively, the production rate *f(y_i_,σ_i_,θ_i_,α_i_)* is raised to *A*(*σ_i_,θ_i_*) if the *y_i_*(*t*) value exceeds a threshold *θ_i_* and lowered to *A*(*σ_i_,θ_i_*)*α_i_* otherwise. *A*(*σ_i_,θ_i_*) is a dampening factor which increases with threshold *θ_i_* and decreases with drug dosage *σ_i_*,. The dynamics are described by the Brownian motion in a double-well potential *U(y_i_,σ_i_,θ_i_,α_i_)*:

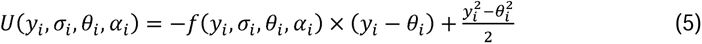

Equation 1 yields two steady pathway activity levels *A*(*σ_i_,θ_i_*) and *α_i_A*(*σ_i_,θ_i_*) corresponding to activation and inactivation of pathway *i*. Hence, we define the “energy gap” Δ*E(σ_i_,θ_i_,α_i_)* between the two states as

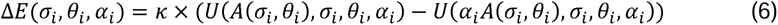

where *k* = 45 for all virtual patients. Transitions between these states are stochastic events and follow a Boltzmann distribution. The transition rate from sensitive to reversible resistant cell states is:

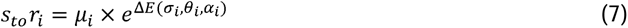

and the transition rate of the opposite direction is:

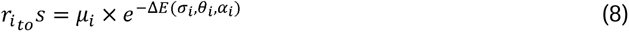

A large energy gap Δ*E(σ_i_,θ_i_,α_i_)* increases the transition probability into the low energy state (inactive pathway or reversible resistant), hence elevates *S_to_r_i_* and lowers *r_ito_ S*.

In the irreversible part of the model, the dynamics of a cellular population also follow a first-order differential equation^21^. The net rate of increase of the cellular population of a phenotype is the sum of the natural net growth rate and the mutation rate into the phenotype, minus the degradation rate induced by drug dosages and the mutation rate out of the phenotype. For instance, denote *n_i_*_0_(*t*) and *n_i_*_1_(*t*) the populations of subclones without and with the irreversible resistance of drug i. Then their population dynamics follow the equations:

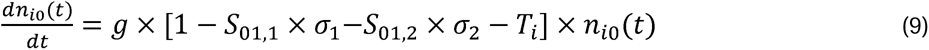

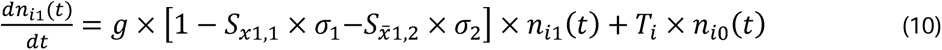

*x* = 1 and *x̄* = 0 for *i* = 1 and *x* = 0 and *x̄* = 1 for *i* = 2. The dynamic equations of the joint model are constructed by combining the reversible and irreversible parts. The net proliferation rate of each cellular population is the balance of production – natural growth and transitions into the designated state through irreversible and reversible resistance mechanisms – and degradation – transitions out of the designated state and reductions by drug dosages. They are illustrated in Fig.1. We also consider the degenerate models of reversible and irreversible resistance alone in Fig. 1. We present an example below, with the full model provided in the supplementary information.

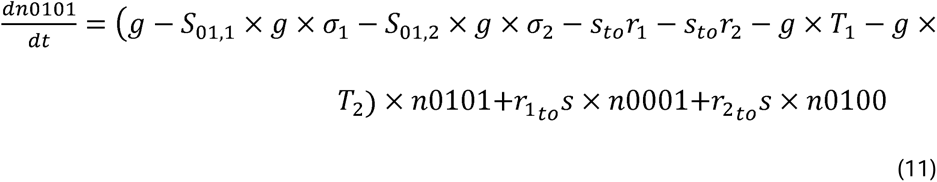

### Treatment strategies

The tumor size and composition in the model are governed by the dynamic equations 11-19 and drug dosage regimens *σ*_1_(*t*) and *σ*_2_(*t*). In principle, *σ_i_*(*t*) can be any temporal function with values in the normalized range [0,1]. In practice, to better mimic the clinical setting of cancer treatment we impose several constraints on *σ_i_*(*t*): (1) 0 ≤ *σ*_1_(*t*) + *σ*_2_(*t*) ≤ 1 V*t*, limiting the total normalized dosage the simulated patient receives at a given time t, as simultaneous full doses of all drugs would often be toxic to the patient. (2) *σ_i_*(*t*) takes possible values of 0, 0.5, 1, reflecting the options to administer either full or half of the drug’s recommended dosage. In clinical settings there are specific maximum doses for each drug individually and for each drug within a simultaneous combination that have been determined from clinical safety data. In these simulations we consider, without loss of generality, the case where each drug is given at half-dose when in simultaneous combinations (3) *σ_i_*(*t*) is a step function which changes values at the beginnings of fixed time intervals – six weeks in our setting, also imitating the periodic administration of drugs, which often are given every 3 weeks. Moreover, radiologic evaluation would generally be no more frequent than every six weeks even in a clinical trial setting.

A treatment strategy is an algorithm for generating *σ_i_*(*t*)’s based on the model parameter values and trajectories of population sizes before the current time point *t*. Ideally, we want to design a treatment sequence to (1) cure the patient (make all cellular populations vanish) if they are curable, and (2) maximize the lifespan of the patient if they are not curable. The optimal solution of *σ_i_*(*t*)’s is difficult to find given the complicated governing equations and restricted temporal functions. Following our previous studies^21–23^ we propose nine heuristic treatment strategies (Fig. 2). These heuristic strategies may be intuitive for oncologists.

**Fig 2:**
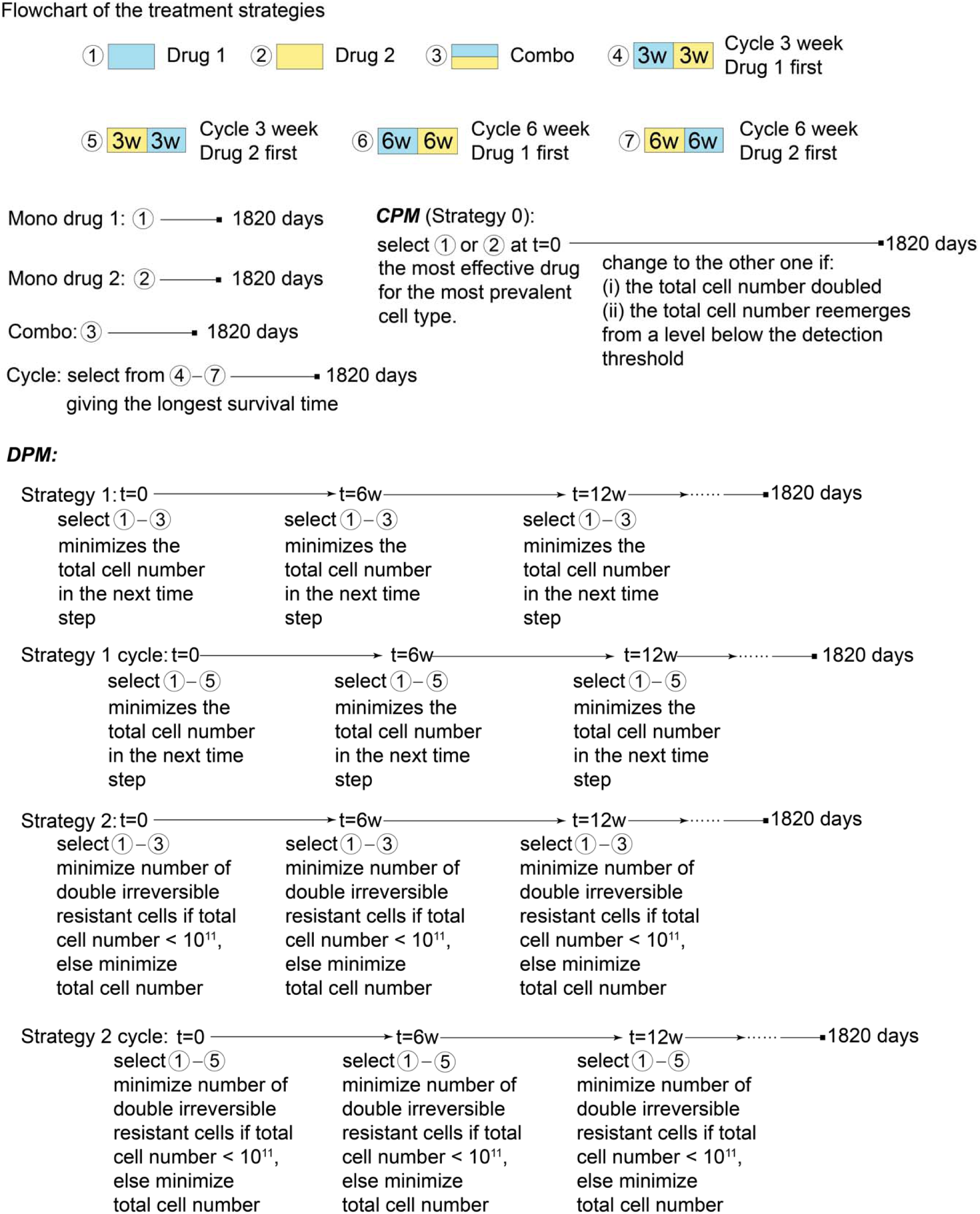
Flowchart of the treatment strategies. Flowchart of the nine treatment strategies outlined in the Methods section.

*Mono drug 1 (Mono1)*: treatment with continuous and constant administration of drug 1. *σ =* (1,0).

*Mono drug 2 (Mono2)*: treatment with continuous and constant administration of drug 2. *σ* = (0,1).

*Combo*: treatment with continuous and constant administration of a combination of half doses of drugs 1 and 2. *σ =* (0.5,0.5).

These three static treatments, Mono1, Mono2 and Combo, are treated as benchmarks against more involved strategies.

*Cycle*: Periodic treatment cycling between drugs 1 and 2. The longest survival time is selected from four possible cycling treatments with combinations of two periods (three or six weeks each) and opposite phases (drug 1 first or drug 2 first). The cyclic treatment is included because it achieves comparable performance to the strictly optimal treatment strategy in our previous model of reversible resistance while being simpler and more practical in clinical application^23^.

*Strategy 0*^21^ *(S0)*: Current personalized medicine (CPM) strategy. Initially, it treats the virtual patient with the most effective drug on the most abundant cellular population. The nadir, a local minimum of the total cell number among the time-series profile, is set equal to the initial total population at t=0, which is 5 ×10^9^, and updated as needed at each time step. The drug is changed if either one of the following events occurs: (i) the total cell number reaches twice the nadir (equivalent to a 25% increase in linear dimensions) or (ii) the total cell number reemerges from a level below the detection threshold (10^9^). If either (i) or (ii) happens and another drug has not been used, switch to another drug which means that each drug is only used once. This results in longer periods before adjustment of the treatment compared to DPM strategies.

*Strategy 1*^21^ *(S1)*: At each time step, select the *σ* from (1,0), (0,1) and (0.5,0.5) that minimizes the predicted total cell number in the next time step. This strategy is intuitive but also myopic.

*Strategy 2*^21^ *(S2)*: Minimize the cell number of the doubly irreversible resistant cell state (1111) unless a meaningful clinical risk is imminent, and switch to minimizing the total cell number if the latter occurs. At each treatment period, select the *σ* from (1,0), (0,1) and (0.5,0.5) that minimizes the predicted cell number of the incurable 1111 state if the total cell number does not exceed 10^11^. This threshold was the most effective threshold of several studied previously in terms of prolonging survival^21^ under the conditions of that simulation. However, this threshold can be adjusted as needed and the optimum may depend on the initial conditions and clinical risk as a function of cell number. In actual clinical cases, many factors would determine whether the treating physician needs to prioritize immediate cytoreduction over prevention of future resistance, and in practice the physician would make this determination. Strategy 2 essentially advises the physician to prioritize resistance prevention unless there is a clear immediate risk that needs to be addressed with immediate cytoreduction. For example, in acute leukemia, immediate cytoreduction in an “induction period” is essential. The use of a simple threshold in this simulation is a necessary approximation based on the great variety of clinical scenarios, but these features can easily be customized in any simulation devoted to a specific application. By preventing the formation of the 1111 cells, the possibility for long-term survival and/or cure is maintained. If the total cell number exceeds 10^11^, select the *σ* from (1,0), (0,1) and (0.5,0.5) that minimizes the predicted total cell number. This strategy attempts to balance the long-term objective of preventing the emergence of incurable tumor cells and the short term objective of shrinking the tumor size.

*Strategy 1 cycle (S1c)*: Same as strategy 1 except adding two cycling treatments as options in each time step. The algorithm selects *σ(t)* from three static treatments – (1,0), (0,1) and (0.5,0.5) – and two cycling treatments – three weeks of drug 1 treatment followed by three weeks of drug 2 treatment in each period, and the cycling treatment with the opposite phase – to fulfill the strategy 1 objective. S1c is a composite strategy combining strategy 1 for the irreversible model and the cyclic treatment for the reversible model but using only a 3 week cycling period.

*Strategy 2 cycle (S2c)*: Same as strategy 2 except adding the aforementioned cycling treatments as options in each time step. S2c is also a composite strategy combining strategy 2 for the irreversible model and the cyclic treatment for the reversible model but using only a 3 week cycling period.

### Model Simulation

We evaluate the effectiveness of the aforementioned nine strategies by conducting simulations on virtual patients with 6131903 distinct parameter configurations. The initial total cell population is 5 × 10^9^, which roughly corresponds to a 5-cm^3^ lesion. The initial cell type composition is determined by parameters R_1ratio_ and R_2ratio_. The time step of predicting responses in each strategy is set to 42 days mimicking the minimum 6-week interval between radiologic evaluations typically on clinical studies^21^. The total simulation period is approximately 5 years, or 1820 days. A cellular population does not proliferate if the size is smaller than 1 cell. A tumor becomes lethal when its population size exceeds 10^13^ cells. This threshold value approximates the total cell numbers in many metastatic lesions, which often lead to mortality quickly^21^. More complex functions providing a monotonically increasing risk of mortality as a function of cell number could be applied if desired. Simulation stops if the total cell number of the virtual patient exceeds 10^13^, or if the virtual patient is cured (the cell number of each state < 1). Therefore, the survival time is either the time when the tumor size reaches the mortal threshold of 10^13^ cells or a maximum of 5 years, indicating that the tumor has either been eliminated or has not caused death within the simulation period. A treatment strategy is significantly better than another one if it shows an absolute improvement of at least 56 days (8 weeks) and a relative improvement of 25% in survival time compared to the other strategy. This is analogous to the typical minimum improvement often considered clinically significant in randomized phase 3 trials in cancer^21^. We implemented the simulation using JetStream2^36^, a resource from the Advanced Cyberinfrastructure Coordination Ecosystem: Services & Support (ACCESS) program^37^.

### Sensitivity analysis

Sensitivity analysis concerns how changes in input parameters affect the outcome of the simulation, which is the survival time. Denote *i*, *j, k* the indices of strategies, parameters, and virtual patients respectively, *p_j_* the value of parameter *j*;, and *ST_ijk_*(*p_j_*) the corresponding survival time. Sensitivity value is defined as^38,39^: *S_i_j_k_ = |ST_ijk_*(*p_j_*) *– ST_ijk_*(*p_j_^−^*)| *+* |*ST_ijk_*(*_Pj_*) *– ST_ijk_*(*p_j_^+^*)|, where *p_j_^−^* and *p_j_^+^* denote the decrement and increment of the parameter value *p_j_*.

## RESULTS

### DPM-based treatments (strategy 1 and 2) outperform the current personalized medicine (strategy 0) treatment in the irreversible resistance only model

We first performed a qualitative check by demonstrating that the simulation outcomes on the degenerate model with irreversible resistance alone were similar to our previous study of DPM^21^. The results are consistent with our previous observation that DPM strategies S1 and S2 outperform the current precision medicine S0 strategy. Fig. 3a displays the Kaplan-Meier curves of the nine treatment strategies on 578884 virtual patients in the irreversible resistance only model. S1 (magenta) and S2 (purple) curves are significantly superior to S0 (green). In Supplementary Table 2, S1 and S2 have higher median survival time, mean survival time, number of cases of survival over 5 years, and number of cases with numerically longer survival time compared to Mono1, Mono2, Combo, Cycle and S0. Supplementary Table 3 shows that the pairwise hazard ratios of S1 and S2 are much lower compared to those of Mono1, Mono2, Combo, Cycle and S0. S2 is slightly better than S1 in terms of the aforementioned metrics, with a median survival time increase of 86 days, a mean survival time increase of 33 days, and an additional 14140 cases surviving over 5 years, resulting in a hazard ratio of 0.935. These results mirror the results in the original simulation of the irreversible model^21^, in which S2 was the superior strategy but had only a slight advantage over S1. Furthermore, by comparing the results between S1 and S1c and between S2 and S2c in Supplementary Tables 2 and 3, we found that adding the cyclic treatment options to DPM strategies only marginally increased survival times (1-2% increase from S1 to S1c and 0.3% increase from S2 to S2c), and yielded pairwise hazard ratios near 1. The limited improvement by incorporating cycling treatment options into DPM is expected because cycling drug dosages are expected to be more effective on reversible resistance cell states, which are absent in the degenerate irreversible resistance only model.

**Fig 3.**
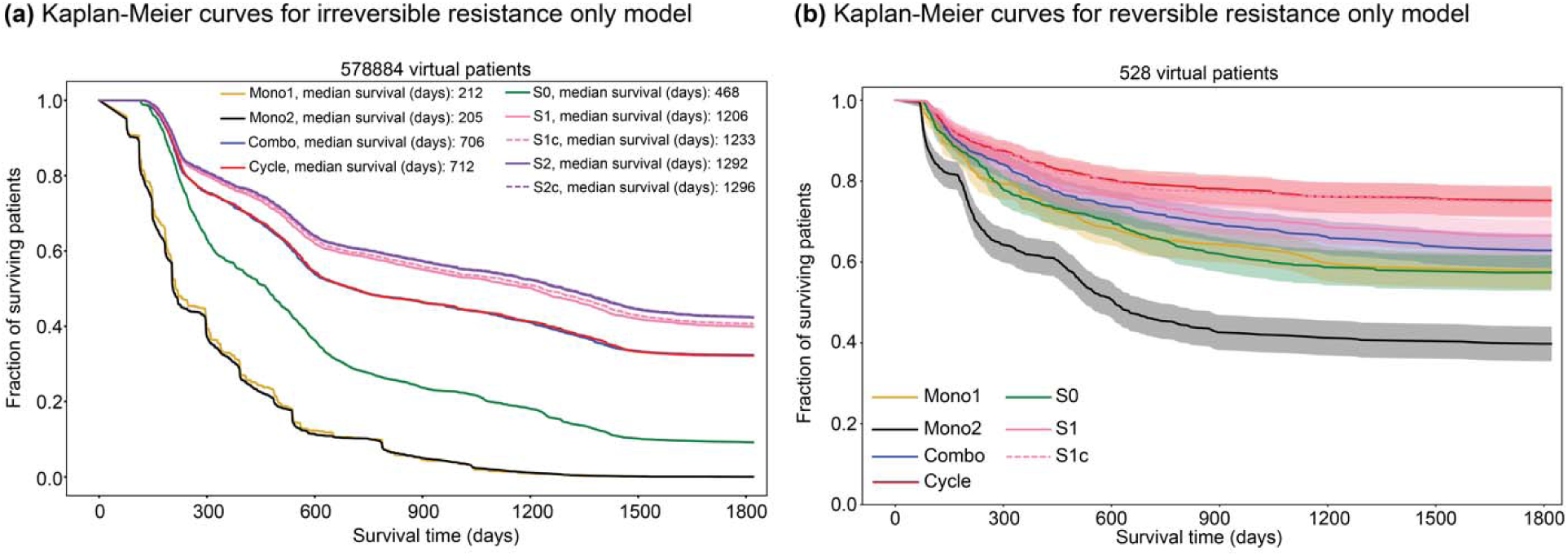
Kaplan-Meier curves of several treatment strategies in the irreversible and reversible resistance only model. **a:** Kaplan-Meier curves of nine treatment strategies over 578884 virtual patients in the irreversible resistance only model. **b:** Kaplan-Meier curves of seven strategies over 528 virtual patients in the reversible resistance only model. The shade of each curve denotes the one standard deviation range of simulations.

Fig. 4a-d illustrates the benefit of the DPM strategy (S2) in a virtual patient possessing a dominant sensitive (0101, ∼90%) cell population and two minor, singly resistant (1101, ∼10% and 0111, <0.001%) cell populations initially. S0 (Fig. 4a) targets the dominant subclone (0101), thus chooses the more effective drug 1 in the first period. Sensitive (0101) and drug 2 resistant (0111) cells are drastically reduced, but drug 1 resistant (1101) cells rapidly proliferate and doubly resistant (1111) cells arise in the first period. Hence the tumor becomes incurable and quickly reaches the mortal threshold. S1 (Fig. 4b) attempts to minimize the predicted total population, thus chooses drug 2 in the first period because it reduces both sensitive (0101) and the abundant drug 1 resistant (1101) cells. The total population is minimized after the first period, but drug 2 resistant cells rapidly proliferate and doubly resistant cells also arise in the first treatment window. Combination of both drugs is administered in the second period to curb the two singly resistant cell populations, but the tumor is still incurable and quickly reaches the mortal threshold. S2 (Fig. 4c) attempts to prevent the emergence of doubly resistant cells, thus chooses two-drug combination in the first period to reduce both singly resistant cell populations. The doubly resistant cells do not arise, hence drug 2 is administered in the subsequent periods to eliminate drug 1 resistant cells. Eventually the patient is cured. Fig. 4d displays the trajectories of total cell numbers of the S0, S1, and S2 strategies. In previous work^21,22^ we have found that initial simultaneous combinations are not always optimal, however; this depends on the detailed dynamics.

**Fig 4.**
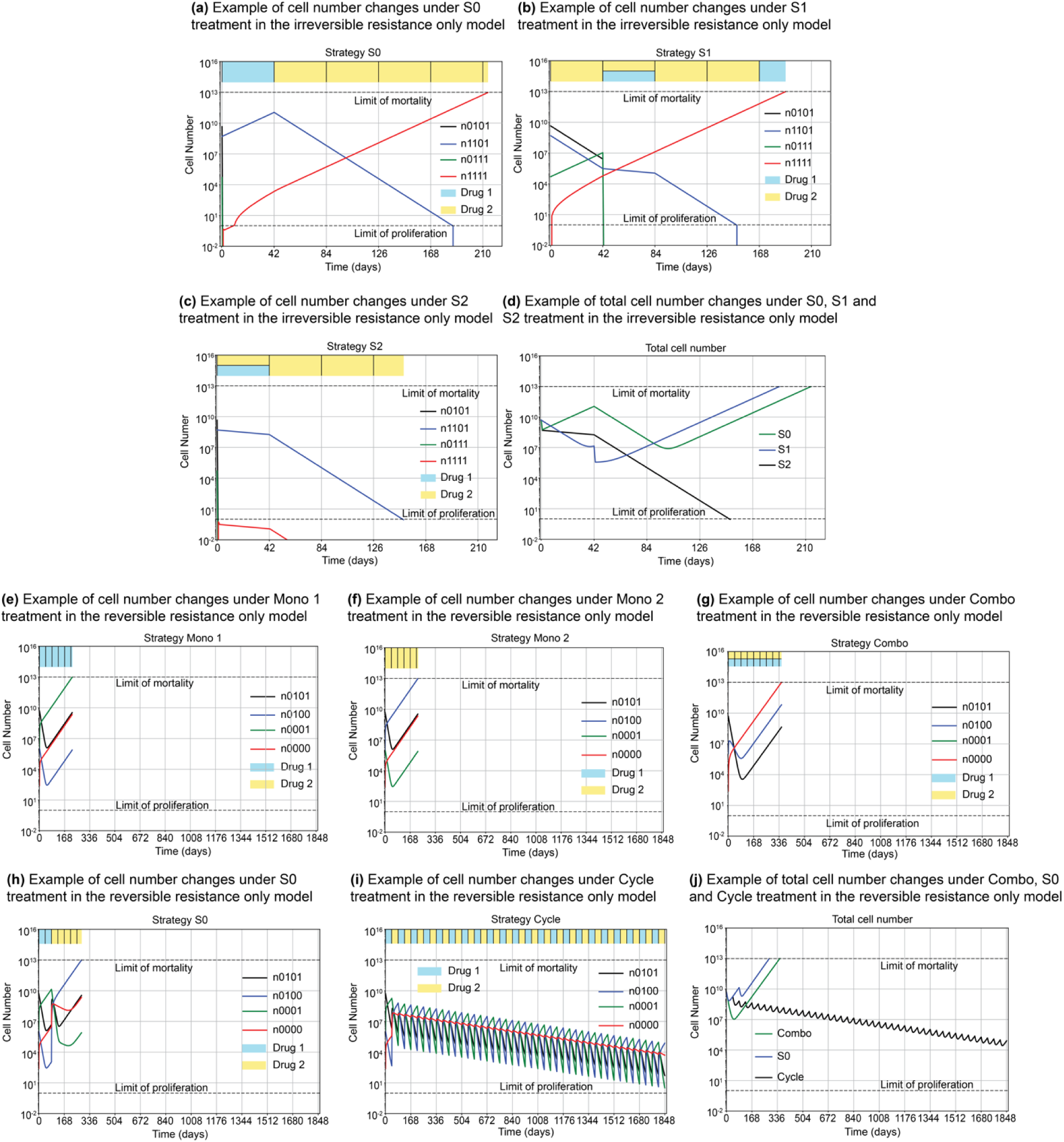
Population dynamics of illustrative virtual patients in the irreversible and reversible resistance only model. **a-d:** Population dynamics of an illustrative virtual patient in the irreversible resistance only model contrasting S0, S1, and S2 treatment responses. Panels **a**-**c** display the population dynamics of individual cell types and dosage sequences under the three treatment strategies. The drug dosage in each treatment period is visualized by colored bars on the top row. Panel **d** displays the population dynamics of the sum of all cell types under the three strategies. **e-j:** Population dynamics of an illustrative virtual patient in the reversible resistance only model contrasting Mono 1, Mono 2, Combo, S0, and Cycle treatment responses. Panel **j** displays the population dynamics of the sum of all cell types under the 5 strategies.

### Cycling treatment outperforms mono and combo treatments in the reversible resistance only model

We performed the second qualitative check by demonstrating that the simulation outcomes on the degenerate model with reversible resistance alone were similar to our previous study of a reversible resistance model^23^. Fig. 3b displays the Kaplan-Meier curves incorporating the seven treatment methods on 528 virtual patients in the reversible resistance only model. The Cycle treatment substantially outperforms other strategies according to both Fig. 3b, survival time metrics in Supplementary Table 4, and pairwise hazard ratios in Supplementary Table 5. Moreover, unlike the irreversible resistance only case, the mean survival time increases about 7% from 1394 days in S1 to 1494 days in S1c, with a hazard ratio of S1c over S1 treatment of 0.725. These improvements corroborate the effectiveness of the cyclic treatments in tackling reversible resistance.

Fig. 4e-j illustrates the effectiveness of the Cycle strategy in a virtual patient possessing a dominant sensitive (0101, ∼99%) cell population, along with two minor singly resistant (0100, ∼0.02% and 0010, ∼0.02%) cell populations, and a smaller minor doubly resistant (0000, ∼0.00005%) cell population initially. As expected, the mono treatment of each drug fails to curb the proliferation of its resistant cells (Fig. 4e and 4f). Simultaneous administration of both drugs (Fig 3i) pushes transitions of the two singly resistant cell populations (0100 and 0001) into the doubly reversible resistant cells (0000), rapidly leading to mortality. S0 (Fig. 4h) also drives transitions into the 0000 cell state and yields an even shorter survival time than Combo (Fig. 4g). By contrast, Cycle (Fig. 4i) pushes cells to oscillate between the two singly resistant states (0100 and 0010) through the intermediate sensitive state (0101). This cycling ensures that there is always a period during which at least one of the drugs can kill the cells, and there is no overlapping period that allows the transition to n0000 cells. Consequently, all cell populations gradually decrease, demonstrating that the Cycle treatment effectively controls the total cell number and outperforms other treatment methods. Fig. 4j displays the trajectories of total cell numbers of the Combo, Cycle and S0 strategies.

### Incorporating cyclic options into DPM-based strategies improve outcomes in the joint model

After confirming the effectiveness of DPM and cycling treatment in the degenerate models of irreversible and reversible resistance, respectively, we sought to demonstrate that the combination of these two approaches can tackle both irreversible and reversible resistance jointly. Fig. 5a displays the Kaplan-Meier curves of nine treatment strategies over 6131903 virtual patients in the joint model. S1c (magenta dashed) and S2c (purple dashed) curves are substantially higher than other strategies. These two strategies also have higher median survival time, mean survival time and number of cases of survival over 5 years compared to the other strategies as shown in Table 2, and smaller pairwise hazard ratios as shown in Table 3. S2c is better on average than S1c based on the metrics on Table 2, with an additional 12312 cases surviving over 5 years. Relative performance of the two strategies varies in individual virtual patients based on the relative potency of the two drugs and other factors. In general, whether S1c or S2c are preferred may need to be individualized in subgroups of patients based on their drug sensitivities and evolutionary dynamics. We note that in our prior work with irreversible model, S1 was also nearly as effective as S2, but S2 had a decisive advantage in some virtual patients^21^. The benefits of adding the cyclic treatment as a treatment option in each period are seen by comparing S1c/S2c with S1/S2 simulation outcomes. S1c/S2c outperforms S1/S2 by 8% in terms of the median survival time, 9% and 7%, respectively, in terms of the mean survival time, and pairwise hazard ratios are approximately 0.92 for each. These improvements in the joint model but not in the irreversible resistance only model, though modest, underscore the effectiveness of cycling treatments in addressing reversible resistance mechanisms when both resistance mechanisms are present. Likewise, the benefits of introducing DPM to tackle irreversible resistance are seen, as the Cycle strategy (cycling drug 1 and 2 without DPM) has very similar outcomes to S1c and S2c in the reversible resistance only model (Supplementary Tables 4 and 5), but S1c is modestly superior to Cycling in the joint model (Tables 2 and 3). Overall, comparing cycling, S1c, and S2c, modest trends in average results are seen as described above. The benefits are more modest because the cancer model incorporates several distinct processes of resistance development, a greater challenge, But the strategy of choice depends on the dynamics in individual virtual patients in a complex way. It is expected that the benefits of a particular strategy will be seen more in subgroups of patients.

**Table 2.**
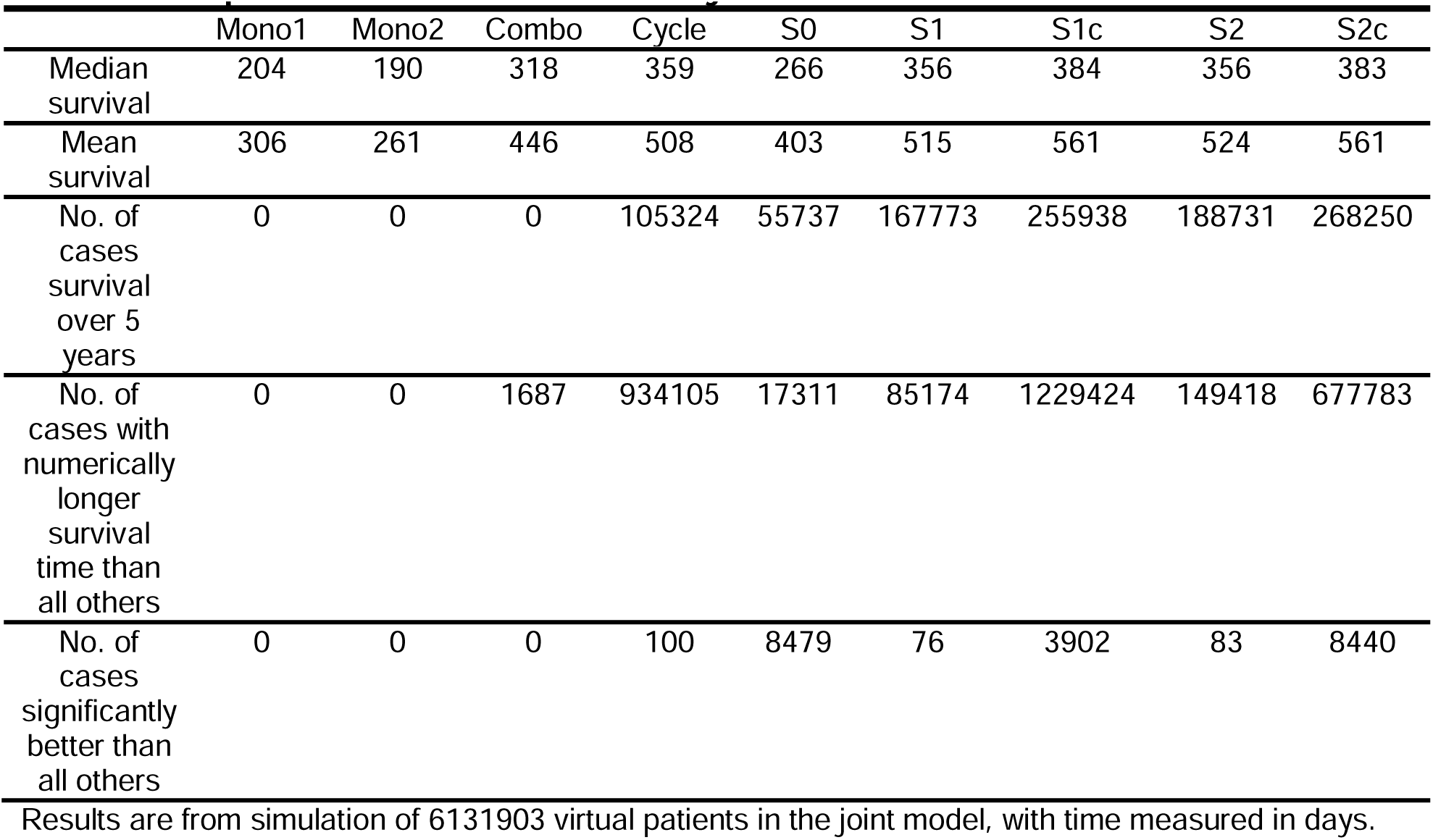
Comparison of treatment results in joint model.

|  | Mono1 | Mono2 | Combo | Cycle | S0 | S1 | S1c | S2 | S2c |
| --- | --- | --- | --- | --- | --- | --- | --- | --- | --- |
| Median survival | 204 | 190 | 318 | 359 | 266 | 356 | 384 | 356 | 383 |
| Mean survival | 306 | 261 | 446 | 508 | 403 | 515 | 561 | 524 | 561 |
| No. of cases survival over 5 years | 0 | 0 | 0 | 105324 | 55737 | 167773 | 255938 | 188731 | 268250 |
| No. of cases with numerically longer survival time than all others | 0 | 0 | 1687 | 934105 | 17311 | 85174 | 1229424 | 149418 | 677783 |
| No. of cases significantly better than all others | 0 | 0 | 0 | 100 | 8479 | 76 | 3902 | 83 | 8440 |

Results are from simulation of 6131903 virtual patients in the joint model, with time measured in days.

**Table 3.** The hazard ratio pairwise comparisons between different strategies in joint model.

|  | Mono1 | Mono2 | Combo | Cycle | S0 | S1 | S1c | S2 | S2c |
| --- | --- | --- | --- | --- | --- | --- | --- | --- | --- |
| Mono1 | N.A. | 0.82 | 1.623 | 1.88 | 1.417 | 1.891 | 2.083 | 1.919 | 2.076 |
| Mono2 | 1.22 | N.A. | 1.928 | 2.212 | 1.699 | 2.228 | 2.437 | 2.259 | 2.429 |
| Combo | 0.616 | 0.519 | N.A. | 1.199 | 0.898 | 1.224 | 1.361 | 1.249 | 1.36 |
| Cycle | 0.532 | 0.452 | 0.834 | N.A. | 0.762 | 1.028 | 1.143 | 1.05 | 1.144 |
| S0 | 0.706 | 0.589 | 1.114 | 1.313 | N.A. | 1.339 | 1.48 | 1.363 | 1.479 |
| S1 | 0.529 | 0.449 | 0.817 | 0.973 | 0.747 | N.A. | 1.11 | 1.021 | 1.111 |
| S1c | 0.48 | 0.41 | 0.735 | 0.875 | 0.676 | 0.901 | N.A. | 0.92 | 1.001 |
| S2 | 0.521 | 0.443 | 0.801 | 0.952 | 0.734 | 0.98 | 1.087 | N.A. | 1.088 |
| S2c | 0.482 | 0.412 | 0.735 | 0.874 | 0.676 | 0.9 | 0.999 | 0.919 | N.A. |

Results are from simulation of 6131903 virtual patients in the joint model. Row strategies are compared to the column strategies.

**Fig 5.**
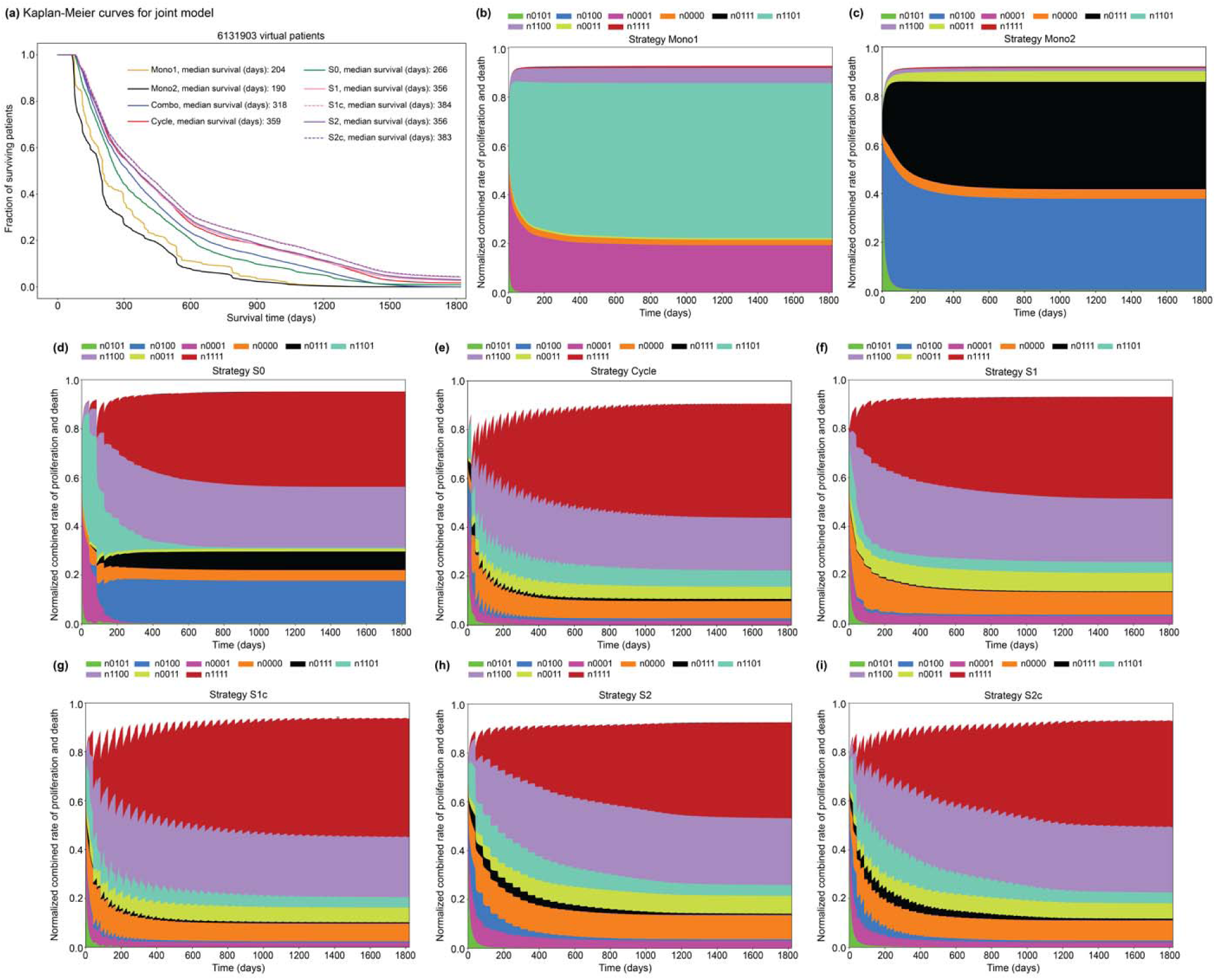
Kaplan-Meier survival curves and rates of each cell state change in the joint model incorporating different treatment strategies and normalized combined rate of proliferation and death for each cell state. 6131903 virtual patients were treated by each of the nine treatment strategies. **a:** Kaplan-Meier curves illustrating the simulated virtual patients. The y axis represents the fraction of surviving virtual patients. Mono1: yellow; Mono2: black; Combo: blue; Cycle: red; S0: green; S1: pink; S1c: dashed pink; S2: purple; S2c: dashed purple. **b-i:** Normalized combined rate of proliferation and death for each cell state across all virtual patients under different treatment strategies. n0101 cells are shown in green, n0100 cells are shown in blue, n0001 cells are shown in purple, n0000 cells are shown in orange, n0111 cells are shown in black, n1101 cells are shown in cyan, n1101 cells are shown in light purple, n0011 cells are shown in yellow and n1111 cells are shown in red. **b:** Mono1 treatment, **c:** Mono2 treatment, **d:** Cycle treatment, **e:** S0 treatment, **f:** S1 treatment, **g:** S1c treatment, **h:** S2 treatment, **i:** S2c treatment.

Beyond survival outcomes, we are also interested in exploring the influence of treatment strategies on cell type composition dynamics. As a crude metric to capture the average composition dynamics, we calculate the net rate of cell number change for each cellular state (proliferation – death) at each time interval, normalized by the rates of all cell states and over all virtual patients. This metric quantifies the proportion of cells arising at each time interval rather than the accumulated cell numbers up to the specific time, and also takes an average over the trajectories among all virtual patients. Fig. 5b-i displays the cell type composition dynamics of eight treatment strategies. All treatment strategies have rather smooth temporal profiles in their composition dynamics (except for the ripples for the strategies incorporating cycling treatments and the slightly more complicated profiles initially for S0). The dominant cell states for Mono1 (Fig. 5b) and Mono2 (Fig. 5c) are singly resistant to drugs 1 (0001 and 1101) and 2 (0100 and 0111) respectively. S0 (Fig. 5d) mainly selects drug 1 initially and then switches to drug 2 without switching back, hence the major cell types at steady state are resistant to drug 2, including 0100, 0000, 0111, 1100, and 1111. For the other five strategies (Fig. 5e-i), the major cell types at steady state are 1111 and 1100. These strategies switch back and forth between drugs 1 and 2, leading to cells that are resistant to both drugs. The relatively small portion of 0011 cells compared to 1100 cells might be due to the high potency of drug 1 compared to drug 2, where cells require mutations of drug 1 to survive. S2 and S2c also have a relatively lower portion of 1111 cells compared to Cycle, S1 and S1c. This is because S2 and S2c are designed to specifically target and minimize the emergence of doubly irreversible resistant cells.

In Fig. 6, we presented an example of a virtual patient where S2c yields the longest survival time. Fig. 6a-b display the dynamics of non-mutated and mutated cell populations for the Cycle strategy. In Fig. 6a, Cycle effectively eliminates non-mutated cells such as 0101, 0100, 0001 and 0000. In Fig. 6b, Cycle cannot eliminate cells that are irreversible resistant to drug 1 in this case. Although drug 2 has the potential to eradicate these cells, but it fails to do so since it is only used with a 50% duty cycle. In Fig. 6c and 6d, the dynamics of cell states changes under S1c are similar to the changes under Cycle, where the emergence of 1100 cells contribute to treatment failure. In S1c, after about four time steps, the strategy also selects cycling between drug 1 and drug 2 to minimize total cell numbers. The cell state changes under S2 and S2c are shown in Fig. 6e-h. The primary difference between S2 or S2c and Cycle or S1c is that S2 and S2c doesn’t fully eliminate non-mutant cells. Instead, they aimed to minimize the number of 1111 cells, which indirectly minimizes the 1100 cell numbers since this cell type, with irreversible resistance to drug 1 and reversible resistance to drug 2, is not readily treatable and will eventually mutate to 1111 cells. In our model, a cell state can only proliferate when its cell number is greater than 1. Therefore, S2 and S2c try to prevent the formation of the first 1100 cell. S2 and S2c initially use the less potent drug 2. S2 treatment has fewer options and fails to prevent the formation of the first 1100 cell. However, S2c can use cycling treatment to successfully avoid the formation of 1100 cells, ultimately providing a cure. It is also notable that the first 2 or 3 time steps might be crucial for the ultimate treatment outcome. Once the first incurable cell emerges, all drug treatment options become ineffective, leading to an eventual collapse.

**Fig 6.**
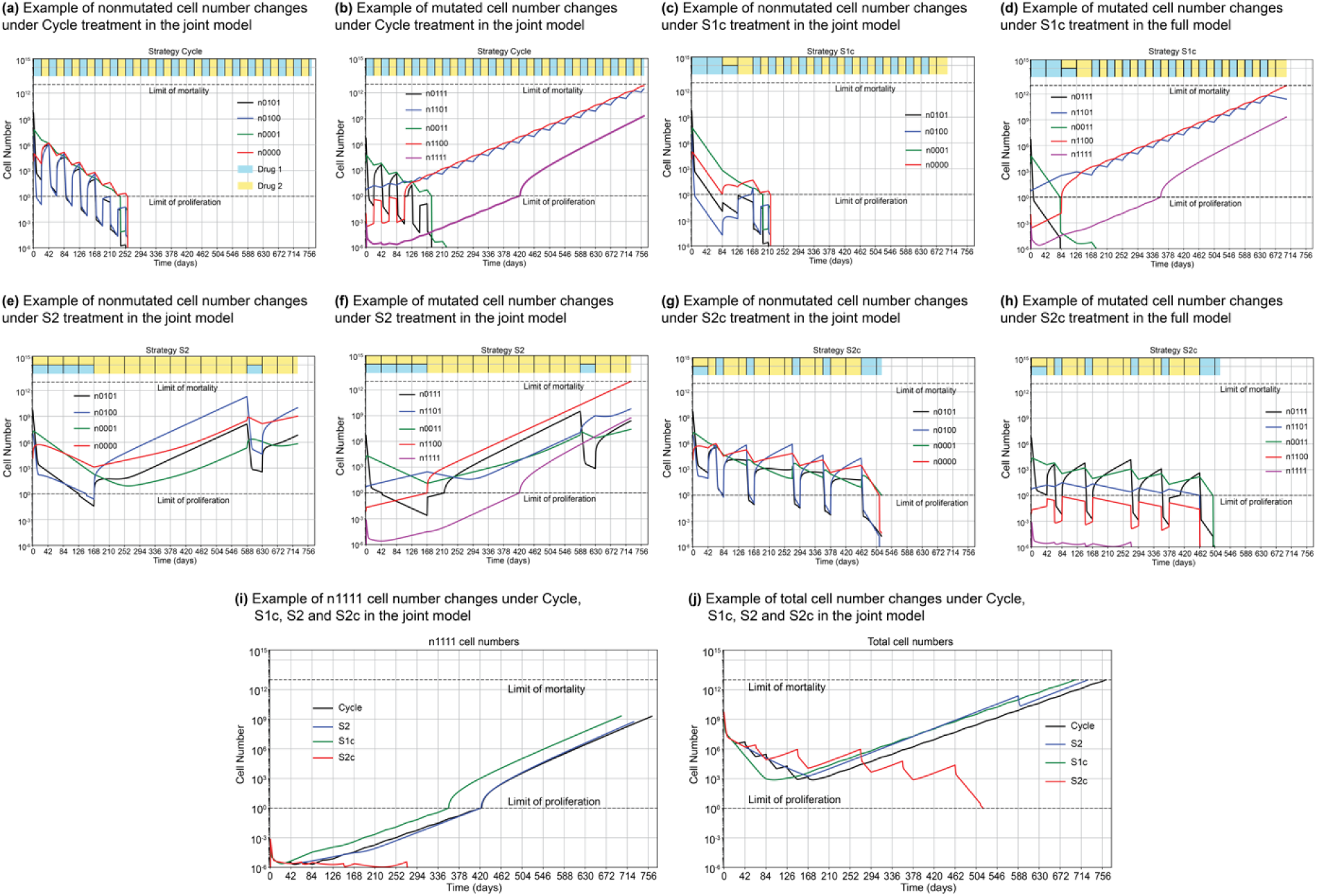
Population dynamics of an illustrative example contrasting four treatment strategies in the joint model. The visual representation in each panel is the same as Fig 2c-j. **a:** dynamics of nonmutated cell numbers (bits 1 and 3 are 0 in their states) under the Cycle strategy. **b:** dynamics of mutated cell numbers (bits 1 or 3 are 1 in their states) under the Cycle strategy. **c:** dynamics of nonmutated cell numbers under S1c. **d:** dynamics of mutated cell numbers under S1c. **e:** dynamics of nonmutated cell numbers under S2. **f:** dynamics of mutated cell numbers under S2. **g:** dynamics of nonmutated cell numbers under S2c. **h:** dynamics of nonmutated cell numbers under S2c. **i:** dynamics of double mutated 1111 cell numbers under four strategies. **j:** dynamics of total cell numbers under four strategies.

### Changes in the treatment strategy for virtual patients when extending the single resistance model to the joint model

Each virtual patient parameter in the joint model can be generated by expanding one corresponding parameter from either of the degenerate models. A virtual patient from the irreversible resistance only model can be extended into virtual patients in the joint model by incorporating parameter α_1_, θ_1_, μ_1_, α_2_, θ_2_, μ_2_, SR_00,1_ and SR_00,2_. Similarly, a virtual patient from the reversible resistance only model can be extended into virtual patients in the joint model by incorporating parameters T_1_, T_2_, SR_11,1_, SR_11,2_, R1_ratio_ and R2_ratio_. For a virtual patient in a degenerate model, there is a numerically best treatment strategy or strategies. In Fig. 7, we used Sankey diagrams to show how the numerically best treatment diverges when extending from a degenerate model to the joint model. S1* represents either S1 or S1c treatment, and S2* represents either S2 or S2c treatment.

**Fig 7.**
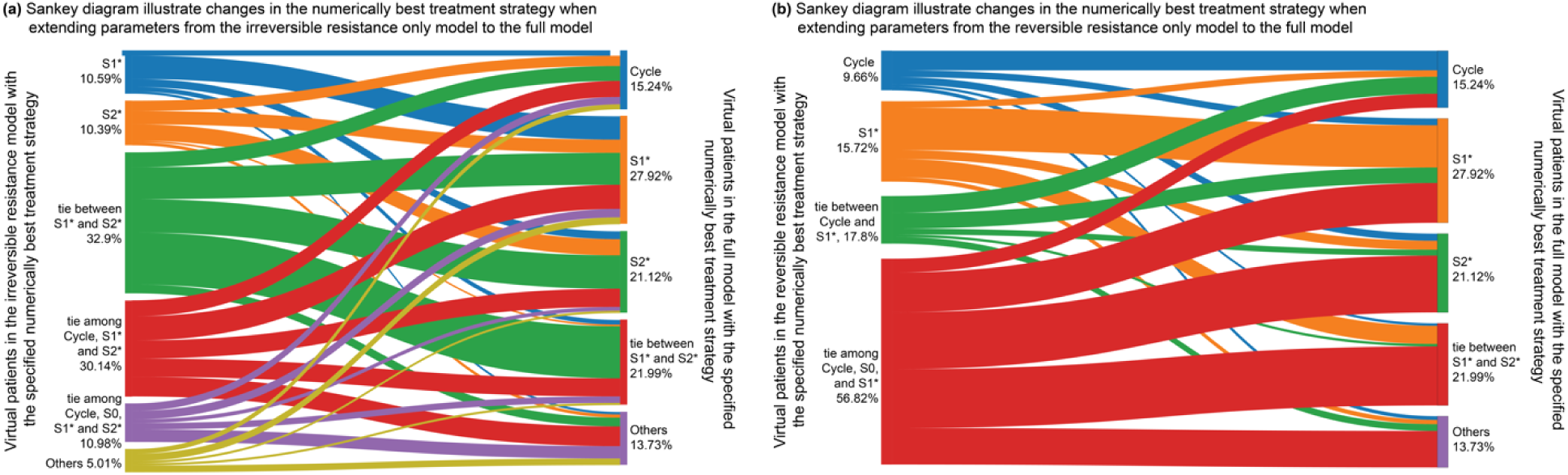
Sankey diagrams illustrate the changes of superior treatment strategy for the virtual patients when extending the degenerate model to the joint model. **a:** Illustrates the changes in the numerically best strategy when extending each virtual patient from the irreversible resistance only model to the joint model. **b:** Illustrates the changes in the numerically best strategy when extending each virtual patient from the reversible resistance only model to the joint model. S1* represents either S1 or S1c treatment. S2* represents either S2 or S2c treatment. Others represents all the other possibilities except those listed in the degenerate model or joint model respectively.

When expanding the virtual patients in the irreversible resistance only model to the joint model, the numerically best treatment strategies diverged as shown in Fig. 7a. For example, of the virtual patients who experienced the longest survival time with the S2* treatment in the irreversible resistance only model, 23.03 % will survive longest under the Cycle treatment in the joint model (see Supplementary Table 6). These changes are due to the incorporation of reversible resistance into the virtual patients and depend on the strength of the resistance. If the introduced reversible resistance is strong, the Cycle treatment is preferred as it can effectively address the reversible resistance. Similarly, when expanding the virtual patients in the reversible resistance only model to the joint model, the numerically best treatment strategies also diverged as shown in Fig. 7b. For example, of the virtual patients who experienced the longest survival time with either the Cycle, S0 or S1* treatment in the reversible resistance only model, 29.62% will survive longest under the S2* treatment in the joint model (see Supplementary Table 7). These changes are due to the incorporation of irreversible resistance into the virtual patients and depend on the strength of the resistance. As the double irreversible resistance cell is incurable, the S2* treatment is preferred as targeting of double resistant cells become crucial.

Overall, the numerically best treatment strategies diverge when expanding from a degenerate model of a single virtual patient to a group of virtual patients in the joint model. The numerically best strategies for individual patients cannot be determined by a single resistance mechanism or predicted solely by the parameters related to one type of resistance in our virtual patient cohort. Instead, they are influenced by the interplay between the two resistance mechanisms, highlighting the necessity of building a unified framework that encompasses both irreversible and reversible drug resistance to identify treatment strategies that effectively address both mechanisms simultaneously.

### Sensitivity analysis of survival time to the parameters in the joint model

To identify which parameters have significant impacts on the outcomes of the nine treatment strategies, we performed a sensitivity analysis of survival time outcomes to the input parameters in the joint model. For each compared parameter or a parameter set (α, θ, μ), we maintained other parameter values constant at the values corresponding to an original virtual patient and calculated the difference of survival times of pairs of virtual patients wherein the one parameter set was incremented or decremented as described in Methods. We consider α, θ, μ, as a single parameter set. If any of these three values differ between two virtual patients, we compared their survival times. For each parameter of T_1_, T_2_, S_01,2_/S_01,1_, SR_11,1_ and SR_11,2_, if the values between two virtual patients differ by one level as detailed in Supplementary Table 1, we compared their survival times. Fig. 8a illustrates the changes in survival time when varying the parameters α, θ and μ, which determine the transition rate between the sensitive and reversible resistant states. These transitions are leveraged by cycling treatments to create opportunities to target and eliminate drug sensitive cells. Monotherapies appears to be less sensitive to these parameters, whereas strategies incorporating cycling treatments are more sensitive. Fig. 8b shows the model’s sensitivity to the ratio of S_01,2_ over S_01,1_, which specifies the efficacy of drug 2 compared to drug 1. Mono 1 is insensitive to drug 2, and all the other strategies have similar sensitivity to this ratio. Fig. 8c illustrates the model’s sensitivity to T_1_ and T_2_. Most the Mono treatments are less sensitive to these transition rates since either the initial resistant subclones will quickly dominate, or the reversible resistance will develop more rapidly. Fig. 8d illustrates the model’s sensitivity to SR_11,1_ and SR_11,2_, the relative sensitivity of the irreversible resistant phenotype to drugs 1 and 2 respectively. Mono 2 is unaffected by SR_11,1_ and Mono 1 is unaffected by SR_11,2_. In addition, survival times are more sensitive to SR_11,1_ than SR_11,2_ since drug 1 is more effective than drug 2. Moreover, the survival times are less sensitive to SR_11,1_ and SR_11,2_ compared to the other parameters. The reduced sensitivity may be due to the relatively small maximum values for SR_11,1_ and SR_11,2_ (8.37 × 10^-2^), which already provide a strong resistance effect. Further decreases in these values have minimal impact on survival time. Overall, the configuration of parameters for each individual virtual patient will influence the outcomes of the treatment strategies, resulting in varying survival times based on different parameter values.

**Fig 8.**
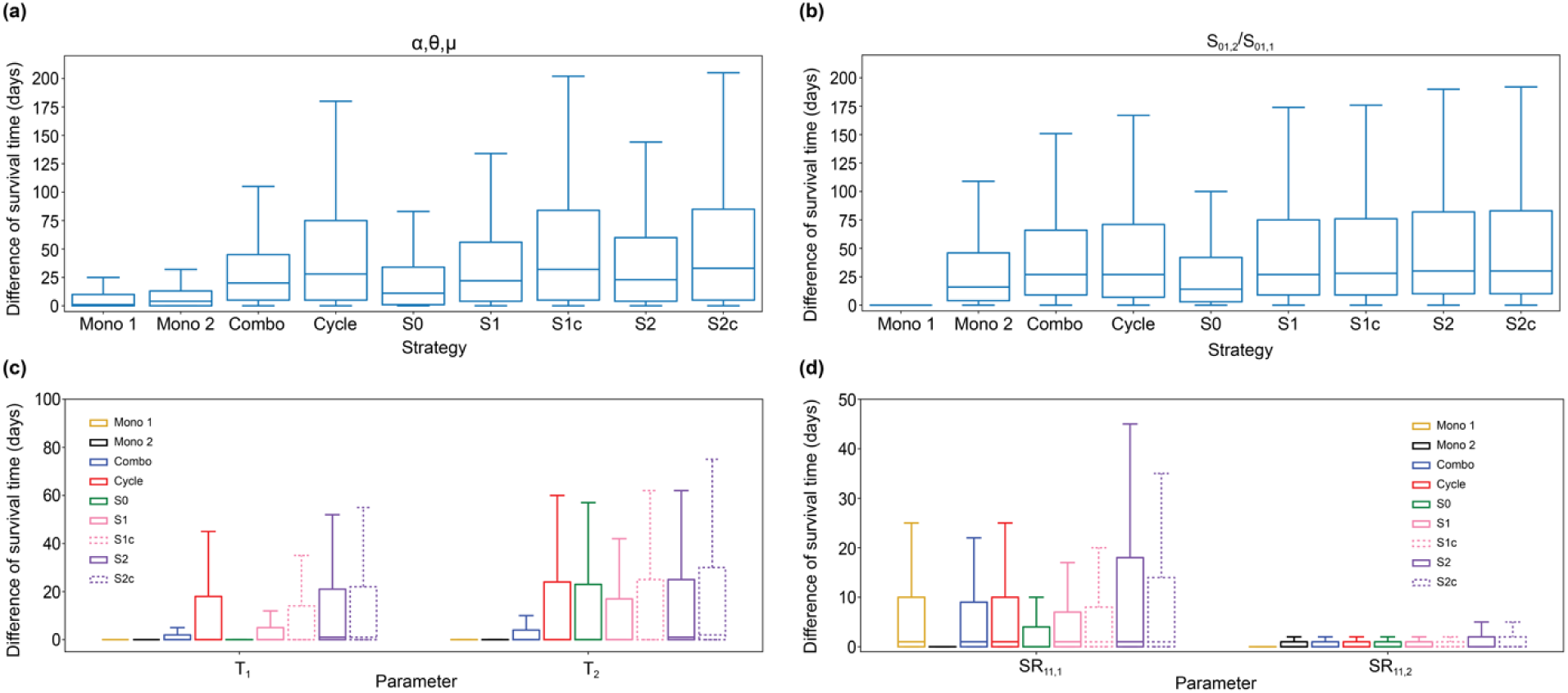
Boxplots illustrating the sensitivity analysis of survival time to the parameters in the joint model. Difference of survival time is on the y axis. **a:** α, θ, μ, (α_1_ equals α_2_, θ_1_ equals θ_2_, μ_1_ equals μ_2_, respectively). **b:** S_01,2_/S_01,1_. **c:** T_1_ and T_2_. **d:** SR_11, 1_ and SR_11, 2_

## DISCUSSION

In this paper, we develop an integrated mathematical model that incorporates both irreversible and reversible resistance mechanisms for two drugs. We evaluate the effectiveness of nine strategies for selecting between 2 possible non-cross resistant treatments by simulating the tumor growth and evolution of six million virtual patients each corresponding to a different set of input parameters. We confirm earlier findings that DPM-based strategies can improve clinical outcomes in a model including only irreversible genetic resistance. We also see an improvement when using only cycling of therapy for non-DPM-based strategies, presumably because this helps with addressing reversible resistance. Our simulation results demonstrate that a combination of the DPM strategy and the cyclic dosages as a treatment option at each period can tackle irreversible and reversible resistance mechanisms simultaneously and hence yields the best performance. The joint treatment strategy takes advantage of the different time scales of reversible and irreversible resistance mechanisms. Knowledge of the reversible resistance kinetics generate optimal cycling building blocks in shorter treatment windows. These short term building blocks are then arranged in an optimal sequence by DPM to address moderate to long term issues.

This work builds upon our previously proposed DPM treatment strategy^21^. DPM is a heuristic and parsimonious model. The model is grounded in oncology principles and, like others, also draws analogies to species evolution. However, species evolution and cancer evolution may be analogous but not identical. Species evolution typically occurs with a lower mutation rate^40^, making the infinite sites approximation that a new mutation at a given base will appear only in one cell at any given time, widely applicable in that context. But this approximation may not hold in cancer where the mutation rate is higher^9,14^. Moreover, cancer evolution is not limited to a single ecological niche, but the niche is continuously expanding and diversifying through multiple metastases and micrometastases, a non-equilibrium situation^31^. Finally, competitive dynamics may differ in micrometastases below the angiogenic limit in size, which can be fully nourished by diffusion^31^. These micrometastases are often part of diffuse organ infiltration and associated with and/or causal of mortality. Moreover, in the high risk neoadjuvant setting, micrometastases may be present below the level of radiologic detection. DPM is also distinguished by its attention to rare subclones and hypermutator subclones. The former accounts for the possible different competitive dynamics, including independent growth in different ecological niches, by not assuming that all rare subclones will be driven to extinction by competition. The latter considers that evolution rate will vary between subclones, a critical aspect of dynamics.

The myopic nature and limitation of the standard personalized medicine approach (S0 in our terminology) for cancer treatment have been criticized and addressed in abundant prior studies, including ours. S0 targets the dominant cell population at each moment, hence often discards the treatment sequences that are suboptimal in the short term but beneficial in the long term. Evolution guided precision medicine models^21–30^, of which DPM is one example, consider intratumoral heterogeneity and dynamics in an attempt to improve upon current precision medicine in a proactive fashion. These approaches have not to date considered the coexistence of drug resistance mechanisms at different time scales in the same framework. Drug resistance phenotypes can be acquired by irreversible mutations or nearly irreversible epigenetic reprogramming, or achieved by reversible switching of the gene regulation/metabolic/signaling pathway activities. Previously, we modeled each type of drug resistance mechanism and provided effective or optimal treatment strategies to tackle them in separate works^21,23^. In this study, we explicitly model these two mechanisms together and provide a simple but effective heuristic to combine them. In the short term (within a treatment period of six-twelve weeks), physicians have options of administering the full dose of single drugs, half dose of both drugs, or alternating the full dose of the two drugs in three week or six week cycles. In the long term (across multiple treatment periods), physicians employ the DPM algorithms (S1 or S2) to design the treatment sequences. DPM may prevent or delay relapse or progression, while cycling may improve response rate and duration. To our knowledge, this is one of the first modeling approaches and treatment strategies for incorporating both irreversible and reversible resistance mechanisms in the same model for two-drug treatment of tumor population dynamics.

The benefits of cycling and of DPM on average remain quite significant compared to the standard precision medicine approach, yet compared to each other cycling and DPM based strategies are more modestly differentiated from each other than seen in the simpler cancer models that considered only reversible resistance or only irreversible resistance. This is expected because the greater complexity of the model creates a greater challenge. Moreover, the optimal strategy is a complex function of the underlying drug properties, heterogeneity, and dynamics. This implies the need to carefully define patient subgroups for whom a given approach will be most optimal, and create evolutionary classifiers to sort individual patients into optimal evolutionary-guided strategies. We have made significant progress in this effort in predicting which patients in the irreversible resistance only model will benefit most from DPM^41^. Experimental validation of our treatment selection strategies will be essential in the future.

Our model should be viewed as a framework that accommodates generic drug resistance mechanisms rather than a realistic model for specific cancer types, targeted drugs, and involved pathways and genes. Nevertheless, there are a number of potential cases that may be captured by this generic modeling framework.

As one example, the availability of several EGFR tyrosine kinase inhibitors (TKIs) for treatment of non-small-cell lung cancer (NSCLC) raises important questions about the optimal treatment sequence^42^. Afatinib, a TKI used for NSCLC, commonly encounters resistance through the T790M gatekeeper mutation, as well as mechanisms involving increased IL6R/JAK/STAT signaling enhanced autophagy^43^. On the other hand, osimertinib as another TKI, is highly selective for treating T790M-positive tumors following failure of prior EGFR TKI treatment and has shown remarkable efficacy in this setting^44^. Resistance mechanisms for osimertinib have also emerged, including *EGFR* mutations in L718V and G724S, HER2 amplification and RAS-MAPK pathway activation^45^. Interestingly, afatinib was successful in overcoming G724S- mediated resistance to osimertinib in vitro^46^, and preclinical studies have shown that L718V mutation retains sensitivity to afatinib^47^. There are reports and clinical studies investigating the feasibility and exploring the optimal sequencing of these two EGFR TKIs^42,48–51^. Our model framework could potentially offer insights into a more effective treatment strategy for these two inhibitors by considering the reversible and irreversible resistance mechanism induced by these EGFR TKIs, with the aim of improving survival outcomes for NSCLC patients.

Another ideal test case for our model framework is triple positive breast cancer (TPBC). TPBC co-expresses two oncogenic drivers, estrogen receptor alpha (ER) and human epidermal growth factor receptor 2 (HER2). Multi-agent adjuvant anti-HER2 and -ER treatment is the superior approach for early stage and advanced or *de novo* metastatic TPBC^52–54^. However, outside academic medical centers a large subset (36.7-60%) of TPBC patients receive single agent anti-ER treatment^55,56^. Re-analysis of a large-scale study of advanced, endocrine-resistant breast cancer^57^ shows that TPBC is overrepresented in those with the highest fraction of their genome altered (q=6.77e-9). These genetic, irreversible resistance mechanisms are accompanied by well-known examples of reversible resistance^58–61^. We anticipate future implementation of this framework will lead to fruitful results in experimental validation and eventually improve treatment outcomes for this and other cancer types.

## DATA AVAILABILITY

This article has no experimental data.

## CODE AVAILABILITY

Code is available at https://github.com/weihevt/DPM-J.

## ACKNOWLEDGEMENTS

This work was funded by the Department of Defense (DoD) Breast Cancer Research Program Awards W81XWH-20-1-0759 and W81XWH-20-1-0760 (to R.B.R and R.A.B., respectively). C.H.Y. was partially funded by a grant (108-2118-M-001-001-MY2) from National Science & Technology Council in the Republic of China (Taiwan). This work used Jetstream2 at Indiana University though allocation MTH240007 from the Advanced Cyberinfrastructure Coordination Ecosystem: Services & Support (ACCESS) program, which is supported by National Science Foundation grants # #2138259, #2138286, #2138307, #2137603, and #2138296.

## AUTHOR CONTRIBUTIONS

W.H., R.B.R., C.H.Y. and R.A.B conceptualized and designed the study. C.H.Y. and W.H. formulated the joint model of drug resistance. W.H. developed the methodology, designed codes, performed simulation, curated data and prepared the initial draft of the manuscript. M.D.M contributed to methodology and data curation. C.H.Y and R.A.B supervised and administered the project. R.B.R. and R.A.B. acquired the financial support for the study. All authors provided critical feedback and helped shape the study and the final manuscript.

## COMPETING INTERESTS

RAB is the Chief Scientific Officer of Onco-Mind, LLC, which owns patents on DPM in the European Union, Taiwan, and Japan. RAB is uncompensated in this role and receives no royalties from these patents.

## Supplementary Information

Equations for the joint model:

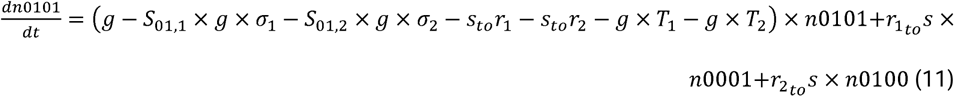

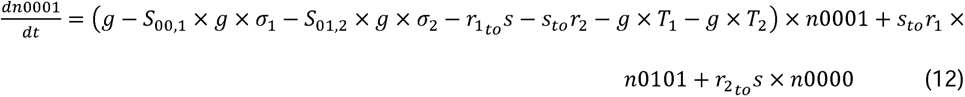

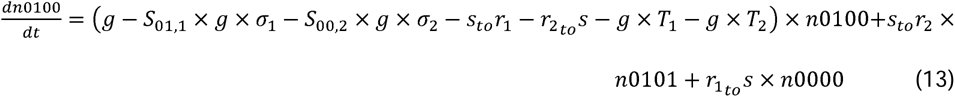

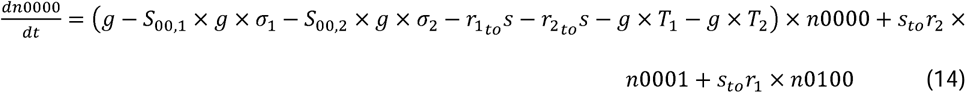

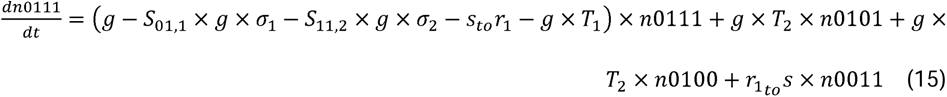

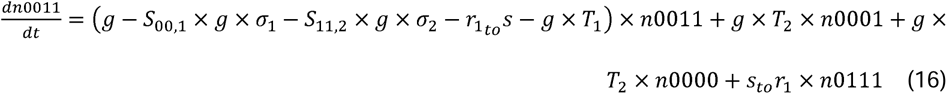

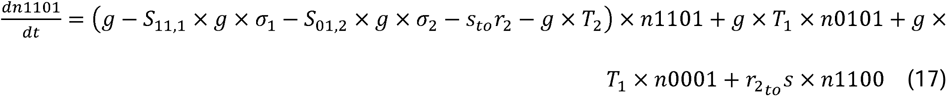

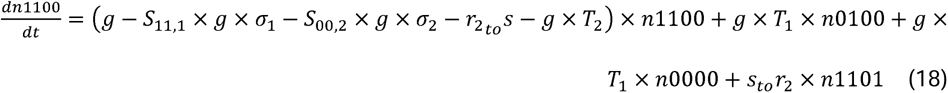

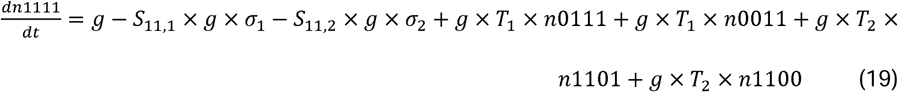

---

**Supplementary Table 1.**
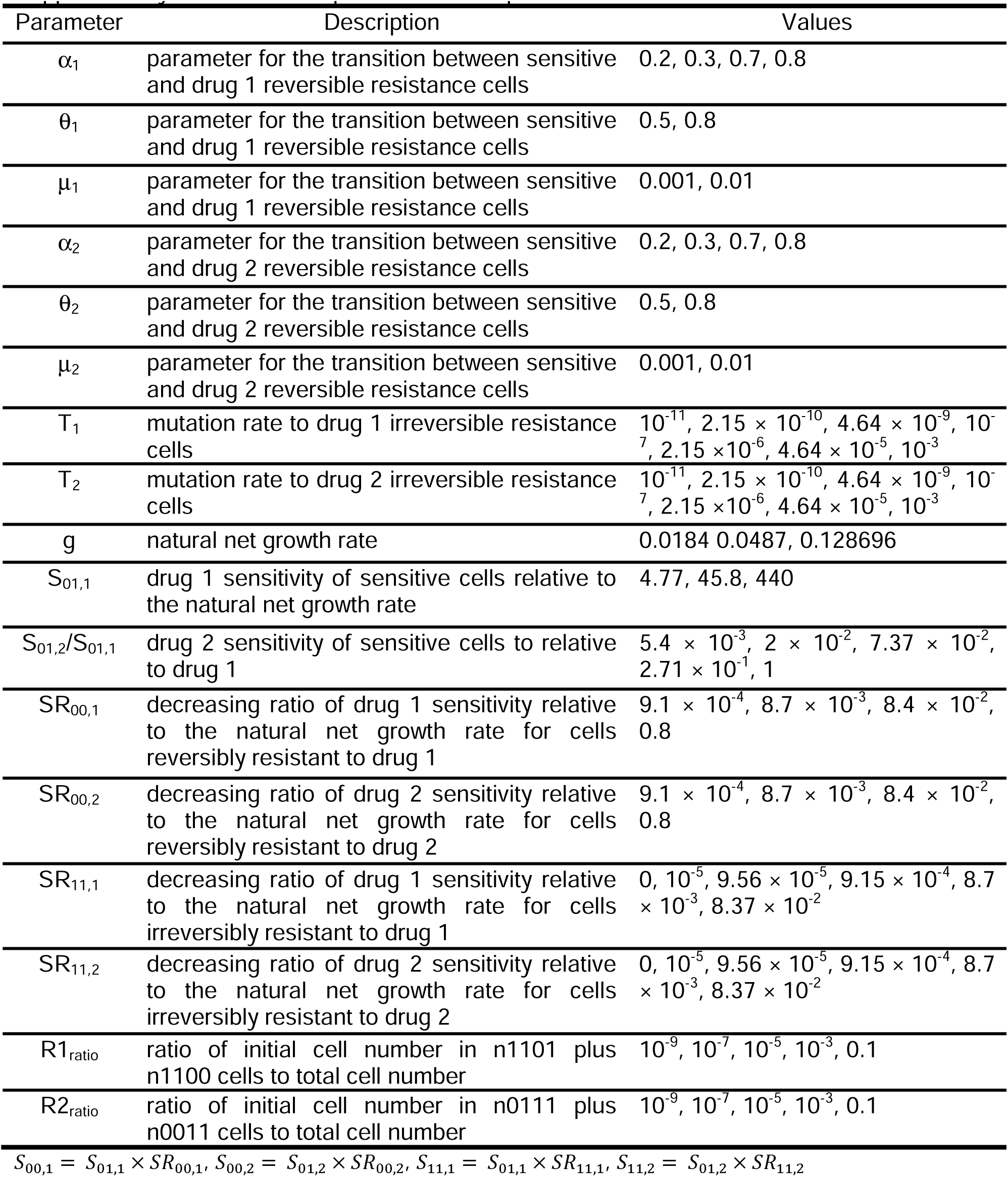
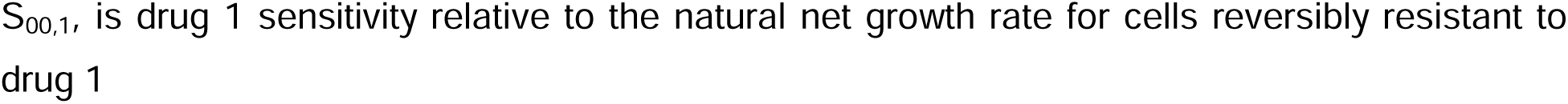
Model parameter description and values.

**Supplementary Table 2.** Comparison of treatment results in irreversible resistance only model.

|  | Mono1 | Mono2 | Combo | Cycle | S0 | S1 | S1c | S2 | S2c |
| --- | --- | --- | --- | --- | --- | --- | --- | --- | --- |
| Median survival | 212 | 205 | 706 | 712 | 468 | 1206 | 1233 | 1292 | 1296 |
| Mean survival | 329 | 321 | 985 | 988 | 643 | 1113 | 1127 | 1146 | 1149 |
| No. of cases survival over 5 years | 154 | 224 | 187383 | 186484 | 53236 | 230878 | 235357 | 245018 | 245904 |
| No. of cases with numerically longer survival time than all others | 0 | 0 | 35 | 1180 | 2090 | 3967 | 31837 | 5869 | 9927 |
| No. of cases significantly better than all others | 0 | 0 | 0 | 0 | 0 | 0 | 0 | 0 | 3 |

Results are from simulation of 578884 virtual patients in the irreversible resistance only model, with time measured in days.

**Supplementary Table 3.** The Hazard ratio pairwise comparisons between different strategies in irreversible resistance only model.

|  | Mono1 | Mono2 | Combo | Cycle | S0 | S1 | S1c | S2 | S2c |
| --- | --- | --- | --- | --- | --- | --- | --- | --- | --- |
| Mono1 | N.A. | 0.963 | 4.083 | 4.129 | 2.305 | 5.268 | 5.44 | 5.637 | 5.672 |
| Mono2 | 1.038 | N.A. | 4.167 | 4.216 | 2.366 | 5.351 | 5.527 | 5.725 | 5.759 |
| Combo | 0.245 | 0.24 | N.A. | 1.005 | 0.526 | 1.246 | 1.28 | 1.332 | 1.339 |
| Cycle | 0.242 | 0.237 | 0.995 | N.A. | 0.522 | 1.242 | 1.275 | 1.328 | 1.334 |
| S0 | 0.434 | 0.423 | 1.902 | 1.915 | N.A. | 2.415 | 2.488 | 2.589 | 2.603 |
| S1 | 0.19 | 0.187 | 0.803 | 0.805 | 0.414 | N.A. | 1.026 | 1.069 | 1.075 |
| S1c | 0.184 | 0.181 | 0.781 | 0.784 | 0.402 | 0.974 | N.A. | 1.042 | 1.047 |
| S2 | 0.177 | 0.175 | 0.751 | 0.753 | 0.386 | 0.935 | 0.96 | N.A. | 1.005 |
| S2c | 0.176 | 0.174 | 0.747 | 0.749 | 0.384 | 0.931 | 0.955 | 0.995 | N.A. |

Results are from simulation of 578884 virtual patients in the irreversible resistance only model. Row strategies are compared to the column strategies.

**Supplementary Table 4.** Comparison of treatment results in reversible resistance only model.

|  | Mono1 | Mono2 | Combo | Cycle | S0 | S1 | S1c |
| --- | --- | --- | --- | --- | --- | --- | --- |
| Median survival, day | 1820 | 608 | 1820 | 1820 | 1820 | 1820 | 1820 |
| Mean survival, day | 1266 | 951 | 1354 | 1499 | 1253 | 1394 | 1494 |
| No. of cases survival over 5 years | 305 | 210 | 332 | 397 | 303 | 351 | 395 |
| No. of cases with numerically longer survival time than all others | 0 | 0 | 0 | 51 | 0 | 2 | 62 |
| No. of cases significantly better than all others | 0 | 0 | 0 | 14 | 0 | 0 | 0 |

Results are from simulation of 528 virtual patients in the reversible resistance only model, with time measured in days.

**Supplementary Table 5.** The hazard ratio pairwise comparisons between different strategies in reversible resistance only model.

|  | Mono1 | Mono2 | Combo | Cycle | S0 | S1 | S1c |
| --- | --- | --- | --- | --- | --- | --- | --- |
| Mono1 | N.A. | 0.58 | 1.201 | 1.893 | 0.992 | 1.356 | 1.867 |
| Mono2 | 1.724 | N.A. | 2.079 | 3.228 | 1.728 | 2.339 | 3.18 |
| Combo | 0.833 | 0.481 | N.A. | 1.584 | 0.823 | 1.129 | 1.56 |
| Cycle | 0.528 | 0.31 | 0.631 | N.A. | 0.523 | 0.714 | 0.985 |
| S0 | 1.008 | 0.579 | 1.216 | 1.912 | N.A. | 1.371 | 1.881 |
| S1 | 0.737 | 0.427 | 0.885 | 1.401 | 0.73 | N.A. | 1.38 |
| S1c | 0.536 | 0.315 | 0.641 | 1.015 | 0.532 | 0.725 | N.A. |

Results are from simulation of 528 virtual patients in the reversible resistance only model. Row strategies are compared to the column strategies.

**Supplementary Table 6.**
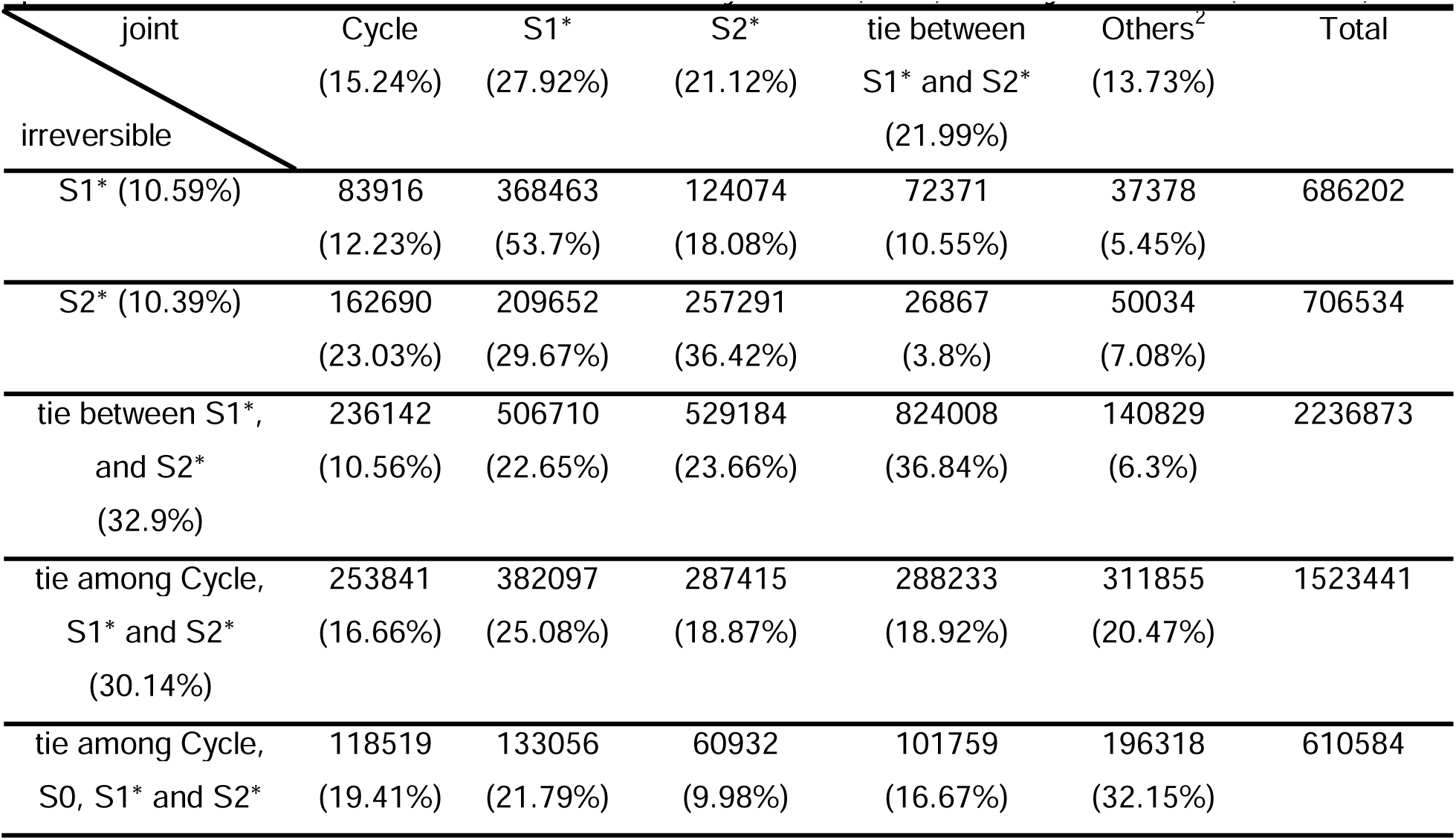
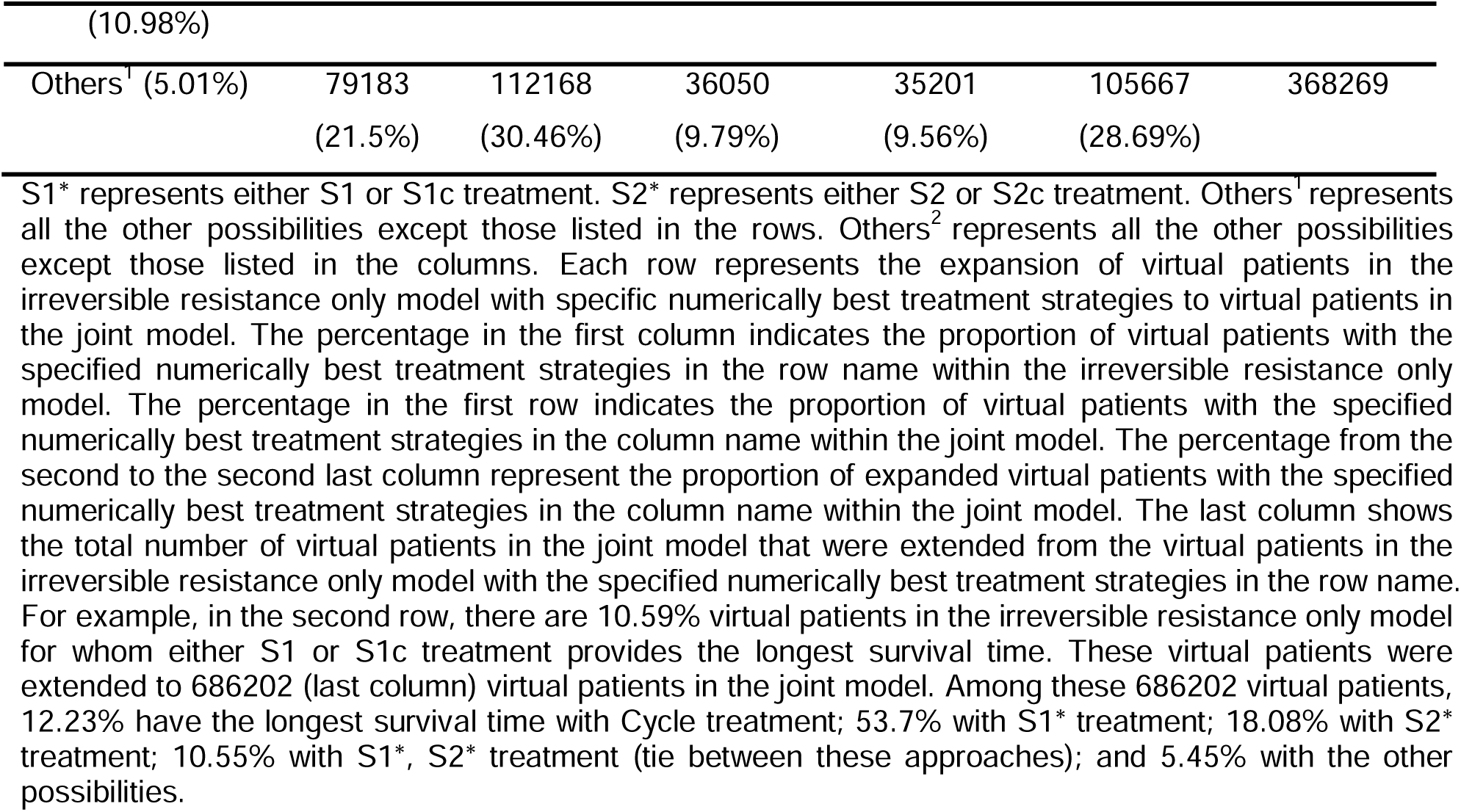
Changes in the numerically best treatment strategy when extending parameters from the irreversible resistance only model (rows) to the joint model (columns)

**Supplementary Table 7.**
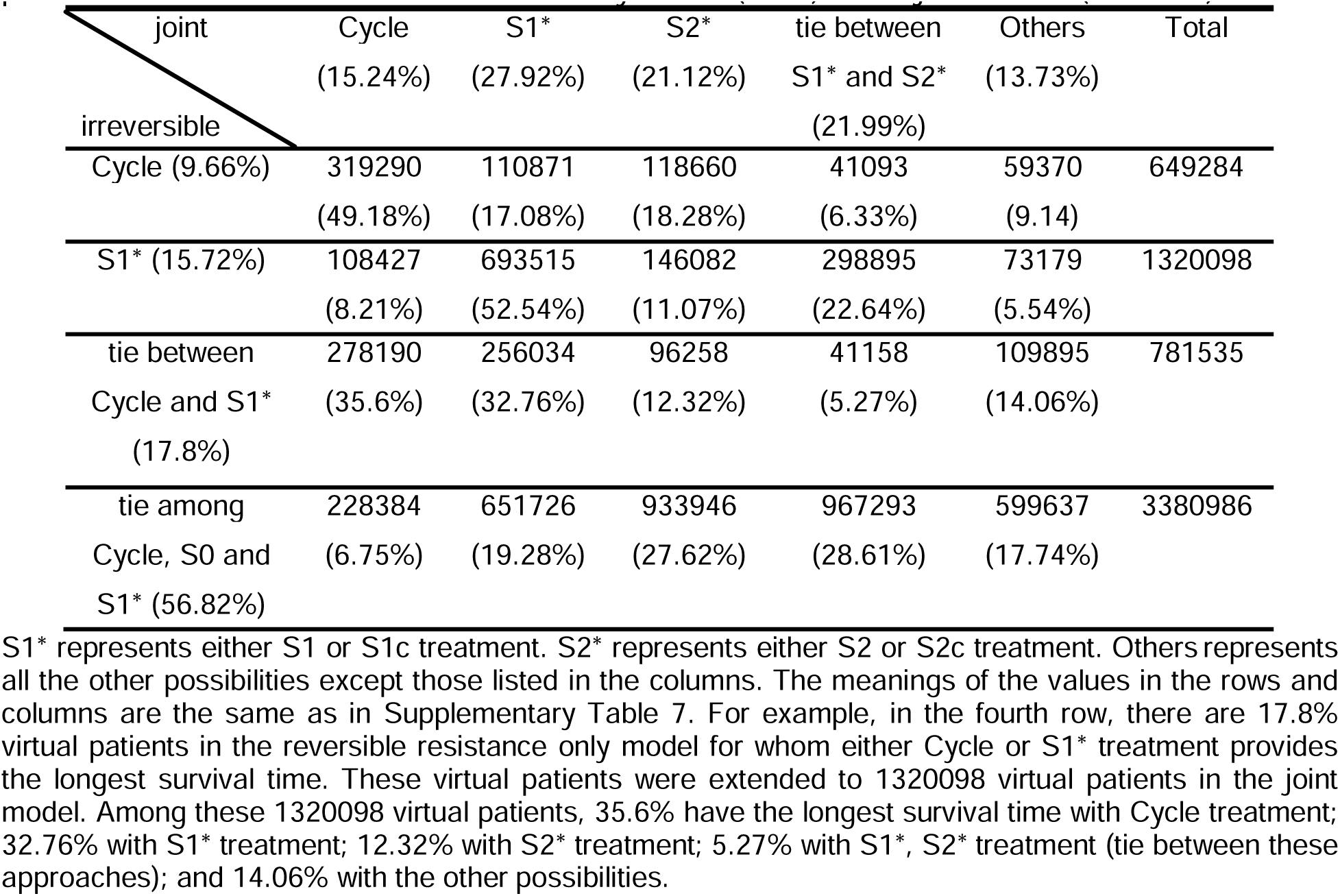
Changes of the numerically best strategy when extending each parameter from the reversible resistance only model (rows) to the joint model (columns)

